# Mitochondrial electron transport chain, ceramide and Coenzyme Q are linked in a pathway that drives insulin resistance in skeletal muscle

**DOI:** 10.1101/2023.03.10.532020

**Authors:** Alexis Diaz-Vegas, Soren Madsen, Kristen C. Cooke, Luke Carroll, Jasmine X. Y. Khor, Nigel Turner, Xin Ying Lim, Miro A. Astore, Jonathan Morris, Anthony Don, Amanda Garfield, Simona Zarini, Karin A. Zemski Berry, Andrew Ryan, Bryan C. Bergman, Joseph T. Brozinick, David E. James, James G. Burchfield

## Abstract

Insulin resistance (IR) is a complex metabolic disorder that underlies several human diseases, including type 2 diabetes and cardiovascular disease. Despite extensive research, the precise mechanisms underlying IR development remain poorly understood. Here, we provide new insights into the mechanistic connections between cellular alterations associated with IR, including increased ceramides, deficiency of coenzyme Q (CoQ), mitochondrial dysfunction, and oxidative stress. We demonstrate that elevated levels of ceramide in the mitochondria of skeletal muscle cells results in CoQ depletion and loss of mitochondrial respiratory chain components, leading to mitochondrial dysfunction and IR. Further, decreasing mitochondrial ceramide levels in vitro and in animal models (under chow and high fat diet) increased CoQ levels and was protective against IR. CoQ supplementation also rescued ceramide-associated IR. Examination of the mitochondrial proteome from human muscle biopsies revealed a strong correlation between the respirasome system and mitochondrial ceramide as key determinants of insulin sensitivity. Our findings highlight the mitochondrial Ceramide-CoQ-respiratory chain nexus as a potential foundation of an IR pathway that may also play a critical role in other conditions associated with ceramide accumulation and mitochondrial dysfunction, such as heart failure, cancer, and aging. These insights may have important clinical implications for the development of novel therapeutic strategies for the treatment of IR and related metabolic disorders.

## Introduction

Insulin is the primary hormone responsible for lowering blood glucose, in part, by stimulating glucose transport into muscle and adipose tissue. This is mediated by the phosphatidylinositol 3-kinase/Akt dependent delivery of insulin sensitive glucose transporters (GLUT4) to the plasma membrane (PM)^1,2^. This process is defective in insulin resistance, a significant risk factor for cardiometabolic diseases such as type 2 diabetes^3^, heart failure^4^, and some types of cancer^5^ and so defective GLUT4 translocation represents one of the hallmarks of insulin resistance.

The development of insulin resistance in skeletal muscle and adipocytes has been associated with multiple intracellular lesions, including mitochondrial Coenzyme Q (CoQ) deficiency^6^, accumulation of intracellular lipids such as ceramides^7^ and increased mitochondrial reactive oxygen species (ROS)^8–10^. However, delineating the relative contribution of these lesions to whole body insulin resistance and their interconnectivity remains a challenge.

Coenzyme Q (CoQ, CoQ9 in rodents and CoQ10 in humans) is a mitochondrial cofactor and antioxidant synthesised and localised in the inner mitochondrial membrane (IMM). This cofactor is essential for mitochondrial respiration^1^^1^, fatty acid oxidation^12^ and nucleotide biosynthesis^13^. We reported that mitochondrial, but not global, CoQ9 depletion is both necessary and sufficient to induce insulin resistance *in vitro* and *in vivo*^6^, suggesting a causal role of CoQ9/10 depletion in insulin resistance. CoQ deficiency can result from primary mutation in the CoQ biosynthetic machinery (named Complex Q)^14^ or secondary from other cellular defects such as deletion of the oxidative phosphorylation system (OXPHOS)^15,16^. Low levels of CoQ10 are associated with human metabolic disease, including diabetes^17,18^, cardiovascular disease^19^ and aging^20^. Strikingly, many of these conditions are also associated with loss of OXPHOS and mitochondrial dysfunction^21,22^. However, it is unclear what causes these mitochondrial defects or if they are mechanistically linked.

Ceramides belong to the sphingolipid family, and high levels are strongly associated with insulin resistance^23^. Whilst it has been proposed that ceramides cause insulin resistance by inhibition of PI3K/Akt signalling^24–26^, there is now considerable evidence that does not support this^3,6,27,28^. Hence, ceramides may induce insulin resistance by a non-canonical mechanism. The rapid decline of mitochondrial oxidative phosphorylation in isolated mitochondria in the presence of N-acetylsphingosine (C2-ceramide), a synthetic ceramide analog that can penetrate cells, suggests that ceramides may be responsible for defective mitochondria^29^. This effect seems to be ceramide-specific as neither diacylglycerides (DAGs) nor triacylglycerides (TAGs) affect mitochondrial respiration ^30^. Recent evidence suggests a link between mitochondrial ceramides and insulin sensitivity, with the observation that reducing mitochondrial, but not global, ceramide in the liver protects against the development of diet induced insulin resistance and obesity^30,31^. Consistent with this, mitochondrial ceramide levels are more strongly associated with insulin resistance than with whole tissue ceramide in human skeletal muscle^30^. Despite this association, no direct evidence exists linking mitochondrial ceramides with insulin sensitivity in skeletal muscle.

Here we describe the linkage between mitochondrial Ceramide, CoQ, OXPHOS and ROS in the aetiology of insulin resistance. We show that a strong inverse relationship between mitochondrial CoQ and ceramide levels is intimately linked to the control of cellular insulin sensitivity. For example, increasing mitochondrial ceramide using either chemical or genetic tools, decreased mitochondrial CoQ levels, and induced insulin resistance. Conversely, genetic or pharmacologic manipulations that lowered mitochondrial ceramide levels increased CoQ levels and protected against insulin resistance. Increased mitochondrial ceramides also led to a reduction in several OXPHOS components, hindering mitochondrial respiration and elevating mitochondrial ROS *in vitro*. This was further supported in human skeletal muscle, where we observed a strong association between insulin sensitivity, abundance of OXPHOS and mitochondrial ceramides. We propose that increased mitochondrial ceramides cause a depletion in various OXPHOS components, leading to mitochondrial malfunction and deficiency in CoQ, resulting in increased ROS and insulin resistance. This provides a significant advance in our understanding of how ceramide causes mitochondrial dysfunction and insulin resistance in mammals.

## Results

### Palmitate induces insulin resistance by increasing ceramides and lowering CoQ9 levels in L6 –myotubes

Lipotoxicity plays a major role in insulin resistance and Cardiometabolic disease^32^. Excess lipids accumulate in insulin target tissues, such as muscle, impairing insulin-stimulated GLUT4 translocation as well as other metabolic actions of insulin. For this reason, several in vitro models have been employed involving incubation of insulin sensitive cell types with lipids such as palmitate to mimic lipotoxicity in vivo^9^. In this study we have used cell surface GLUT4-HA abundance as the main readout of insulin response. As shown (Fig. 1A), incubation of L6 myotubes with palmitate (150 μM for 16 h) reduced the insulin-stimulated translocation of GLUT4 to the cell surface by ∼30 %, consistent with impaired insulin action. Despite this marked defect in GLUT4 translocation, we did not observe any defect in proximal insulin signalling as measured by phosphorylation of Akt or TBC1D4 (Fig. 1C & D - 100nM insulin; 20mins). Previous studies linking ceramides to defective insulin signalling have utilised the short chain ceramide analogue (C2-ceramide)^24–26^. Intriguingly, we were able to replicate that C2-ceramide inhibited both GLUT4 translocation and Akt phosphorylation in L6 myocytes (Fig. 1B, C & D). One possibility is that palmitate induces insulin resistance in L6 myotubes via a ceramide-independent pathway. However, this is unlikely as palmitate-induced insulin resistance was prevented by the ceramide biosynthesis inhibitor myriocin (Fig. 1A) and we observed a specific increase in C16-ceramide levels in L6 cells following incubation with palmitate, which was also prevented by myriocin (Fig. 1E & F, Sup. Fig. 1). Based on these data we surmise that C2-ceramide does not faithfully recapitulate physiological insulin resistance, in contrast to that seen with incubation with palmitate.

**Figure 1.**
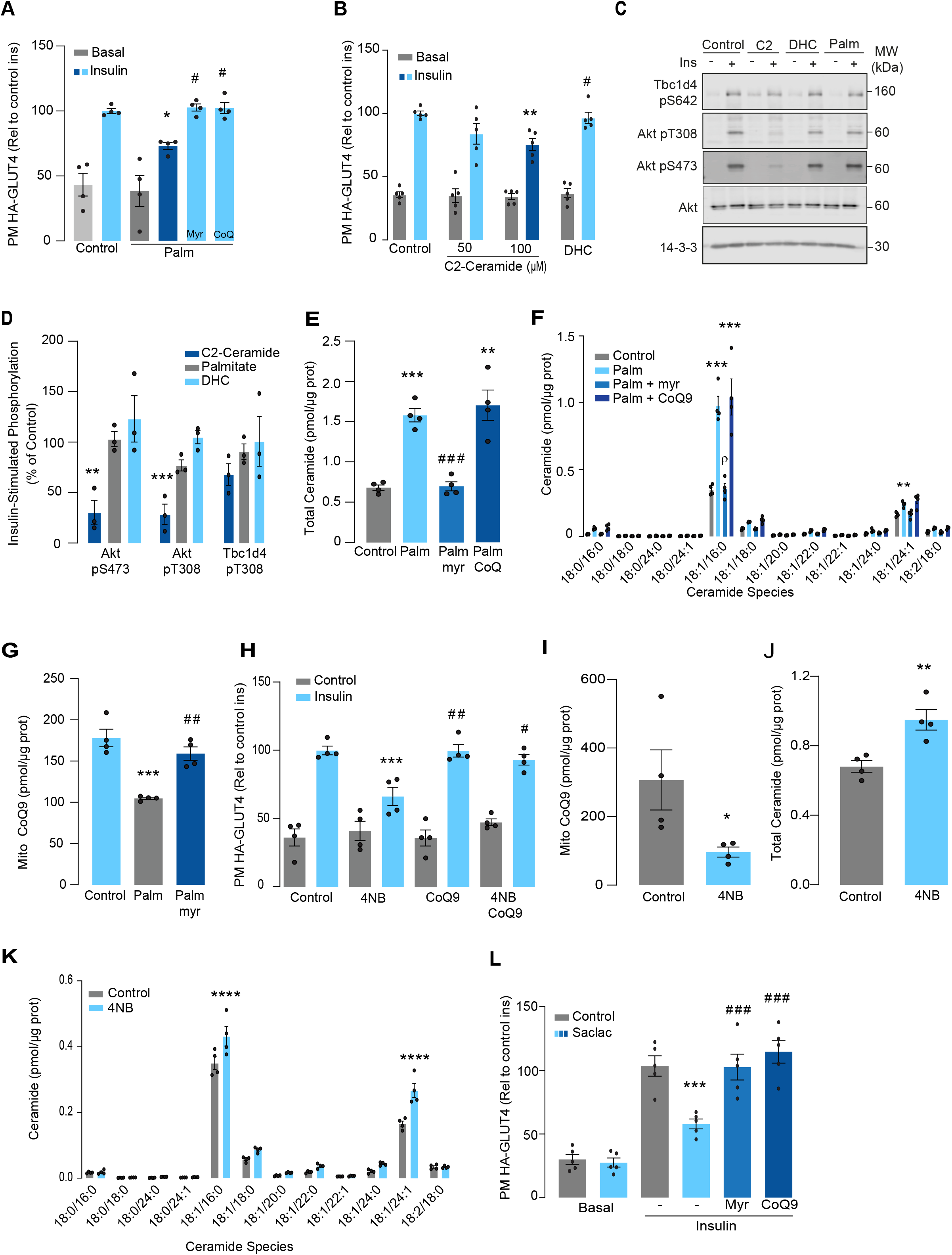
Palmitate increases ceramides, decreases CoQ and induces insulin resistance in L6-myotubes. A) Insulin-induced GLUT4 translocation in L6-HA-GLUT4 myotubes exposed to palmitate (150 μM for 16 h, Palm) or BSA (Control) in presence of DMSO (control), myriocin (10 μM for 16 h) or CoQ9 (10 μM for 16 h). Plasma membrane GLUT4 (PM-GLUT4) abundance was normalised to insulin-treated control cells. N = 4, mean ± S.E.M. *p< 0.05 vs Control ins, # p< 0.5 vs Palm ins B) Insulin-induced GLUT4 translocation in L6-HA-GLUT4 myotubes exposed to C2-ceramide, Dihydroceramide (100 μM, DHC) or DMSO (Control) for 2 h. Plasma membrane GLUT4 (PM-GLUT4) abundance was normalised to insulin-treated control cells. N = 5, mean ± S.E.M. **p< 0.01vs control ins, # p< 0.5 vs 100 μM C2 Ceramide ins. (C and D) L6-HA-GLUT4 myotubes were serum-starved after BSA (Control for 16 h), Palmitate (Palm for 16 h), C2-ceramide (100 μM for 2 h, C2) or dihydroceramide (100 μM for 2 h, DHC) treatment and acute insulin (Ins) was added where indicated. Phosphorylation status of indicated sites was assessed by immunoblot (C). Immunoblots were quantified by densitometry and normalised to insulin-treated control cells (indicated by dotted line). N = 3, mean ± S.E.M. **p< 0.01, ***p< 0.001 (E and F) Endogenous ceramides levels in L6-HA-GLUT4 myotubes treated for 16 h with BSA (control), palmitate (150 μM, Palm), myriocin (10 μM for 16 h) or CoQ (10 μM for 16 h) as indicated in the graph. Total (E) and specific (F) ceramide species were quantified. N = 4, mean ± S.E.M. **p< 0.01, ****p< 0.001vs Control, ### p < 0.001 vs Palm, ρ p<0.01 vs Palm. G) CoQ9 level in mitochondrial fraction obtained from L6-HA-GLUT4 myotubes. N = 4, mean ± S.E.M. ***p< 0.001 vs Control, ## p<0.01 vs Palm H) Insulin-induced GLUT4 translocation in L6-HA-GLUT4 myotubes exposed to 4-NB (2.5 mM for 16 h) or DMSO (Control) in presence of CoQ9 (10 μM for 16 h). Plasma membrane GLUT4 (PM-GLUT4) abundance was normalised to insulin-treated control cells. N = 4, mean ± S.E.M. ***p< 0.001 vs Control ins, ## p<0.01, ### p<0.001 vs 4NB I) CoQ9 level in mitochondrial fraction obtained from L6-HA-GLUT4 myotubes exposed to DMSO (Control) or 4NB for 16 h. N = 4, mean ± S.E.M. *p< 0.05 (J and K) Total (J) and specific (K) ceramide species quantified in L6-HA-GLUT4 myotubes treated for 16 h with DMSO (control) or 4NB (2.5 mM for 16 h).. N = 4, mean ± S.E.M. **p< 0.01, ****p< 0.0001 L) Insulin-induced GLUT4 translocation in L6-HA-GLUT4 myotubes exposed to Saclac (10 μM for 24 h) or EtOH (Control) in presence of DMSO (control), myriocin (10 μM for 16 h) or CoQ9 (10 μM for 16 h). Plasma membrane GLUT4 (PM-GLUT4) abundance was normalised to insulin-treated control cells. N = 5, mean ± S.E.M. ***p< 0.001 vs Control Ins, ### p < 0.001 vs Saclac Ins.

We previously demonstrated that insulin resistance was associated with CoQ depletion in muscle from high-fat diet fed mice^6^. To test if CoQ supplementation reversed palmitate-induced insulin resistance, L6-myotubes were co-treated with palmitate plus CoQ9. Addition of CoQ9 had no effect on control cells but overcame insulin resistance in palmitate treated cells (Fig. 1A). Notably, the protective effect of CoQ9 appears to be downstream of ceramide accumulation, as it had no impact on palmitate-induced ceramide accumulation (Fig. 1E-F). Strikingly, both myriocin and CoQ9 reversed insulin resistance, suggesting that there might be an interaction between ceramides and CoQ in the induction of insulin resistance with palmitate in these cells. Moreover, we have previously shown that mitochondrial CoQ is a key determinant of insulin resistance^6^ suggesting that ceramides and CoQ may interact in mitochondria. To explore this link, we next examined the effect of palmitate on mitochondrial CoQ levels. As shown (Fig. 1G), palmitate lowered mitochondrial CoQ9 abundance by ∼40 %, and this was prevented with myriocin. To test whether CoQ depletion is downstream of ceramide accumulation, we exposed GLUT4-HA-L6 myotubes to 4-nitrobenzoic acid (4NB) to competitively inhibit 4-hydroxybenzoate:prolyprenyl transferase (Coq2), a limiting step in CoQ9 synthesis^33^. 4NB (2.5 mM for 16 h) decreased mitochondrial CoQ9 to a similar extent as observed in palmitate-treated myocytes (Fig. 1 I) and generated insulin resistance in GLUT4-HA-L6 myotubes (Fig. 1 H). Notably, 4NB mediated insulin resistance was prevented by provision of CoQ9, as previously described^6^. Interestingly, total ceramide abundance was increased in 4NB treated cells albeit to a lesser extent than observed with palmitate, without affecting other lipid species (Fig. 1 J & K, Sup. Fig. 1). One possibility is that CoQ directly controls ceramide turnover^34^. An alternate possibility is that CoQ inside mitochondria is necessary for fatty acid oxidation^12^ and CoQ depletion triggers lipid overload in the cytoplasm promoting ceramide production^35^. In fact, increased fatty acid oxidation is protective against insulin resistance in several model organisms ^36–38^. Future studies are required to determine how CoQ depletion promotes Cer accumulation. Regardless, these data indicate that ceramide and CoQ have a central role in regulating cellular insulin sensitivity.

Since palmitate treatment can have a number of effects beyond ceramides, we next attempted to increase intracellular ceramides by inhibiting the ceramide degradation pathway. We exposed L6 myotubes to different concentrations of Saclac, an inhibitor of acid ceramidase (Kao et al., 2019), for 24 h. Saclac increases ceramides in L6 cells in a dose-dependent fashion, with the largest effect on C16:0 ceramides (Sup. Fig. 1F). Interestingly, Saclac also promoted accumulation of DAGs, sphingosine-1 phosphate (S1P) and sphingosine (SPH), demonstrating the tight interaction between these lipid species (Sup. Fig. 1 B, C & E). Consistent with a role of ceramide in insulin sensitivity, Saclac (10 μM for 24 h) reduced insulin stimulated GLUT4 translocation by 40% (Fig. 1L, vs Control; p<0.001) and this was prevented by myriocin or CoQ9 supplementation (Fig. 1L). Notably, no detectable defects in Akt phosphorylation were observed (Sup. Fig. 1G & H).

To explore if ceramides promote CoQ depletion beyond skeletal muscle, human cervical cancer cells (HeLa) were exposed to Saclac, as previously described (2 μM for 24 h)^39^. Consistent with our observation in L6-myotubes, Saclac increased total ceramide levels (∼6 fold over basal) (Sup. Fig. 2 A & B) and lowered CoQ levels inside mitochondria (Sup. Fig 2C). Of interest, myriocin prevented Saclac-induced-CoQ depletion demonstrating that there is a similar interaction between ceramide and CoQ levels in this human cell line as observed in L6 cells (Sup. Fig. 2D). Moreover, this was relatively specific to CoQ as we did not observe any change in mitochondrial mass with Saclac (Sup. Fig 2 E-G), cell viability (Sup. Fig. 2H) or DAGs abundance (Sup. Fig. 2 I) Regardless, these data indicate that there is a strong association between ceramide and CoQ and that this has a central role in regulating cellular insulin sensitivity.

### Mitochondrial ceramide promotes insulin resistance by lowering CoQ levels

Although mitochondrial ceramides have been linked with insulin resistance in human skeletal muscle^30^, to date, there is no direct evidence linking mitochondrial ceramides with insulin sensitivity. We wanted to determine if ceramide accumulation specifically in mitochondria is associated with altered CoQ levels and insulin resistance. To achieve this, we employed doxycycline-Tet-On inducible^40^ overexpression of a mitochondrial-targeted Sphingomyelin Phosphodiesterase 5 (mtSMPD5) in GLUT4-HA-L6 cells (GLUT4-HA-L6-mtSMPD5) (Fig. 2A). SMPD5 is a murine mitochondria-associated enzyme^41^ that hydrolyses sphingomyelin to produce ceramides^42^. Thus, overexpressing mtSMPD5 should specifically increase ceramides within mitochondria and avoid potential non-specific effects associated with small molecule inhibitors. As expected, doxycycline induced mitochondrial expression of mtSMPD5, as demonstrated by enrichment of SMPD5 in mitochondria isolated from L6 cells (Fig. 2B) and this was associated with increased total mitochondrial ceramides to the same extent as observed with palmitate treatment (Fig. 2C), with the largest increase in C16-ceramide (Fig. 2D). Importantly, mtSMPD5 overexpression did not affect ceramide abundance in the whole cell lysate nor other lipid species inside mitochondria such as cardiolipin, cholesterol and DAGs (Sup. Fig. 3 A, D-J). Intriguingly, mtSMD5 did not affect sphingomyelin levels in mitochondria (Sup. Fig. 3G), consistent with exchange between mitochondrial and extra-mitochondrial sphingomyelin pools to compensate for the degradation induced by SMPD5 overexpression^43^. Consistent with our hypothesis, mtSMPD5 was sufficient to promote insulin resistance in response to submaximal and maximal insulin doses (Fig. 2E). Furthermore, mtSMPD5 overexpression promoted insulin resistance without affecting Akt phosphorylation (Fig. 2F - H), and no differences in total GLUT4 levels were observed across the treatments (Fig. 2F & Sup. Fig. 3B). We next explored if mitochondrial ceramide-induced insulin resistance was mediated by lowering CoQ within mitochondria. In line with our previous results, Mitochondrial CoQ levels were depleted in both palmitate-treated and mtSMPD5-overexpressing cells without any additive effects. This suggests that these strategies to increase ceramides share a common mechanism for inducing CoQ depletion in L6 myotubes (Fig. 2I). Importantly, CoQ9 supplementation prevented both palmitate- and mtSMPD5 induced-insulin resistance (Fig. 2J), suggesting that CoQ depletion is an essential mediator of insulin resistance.

**Figure 2.**
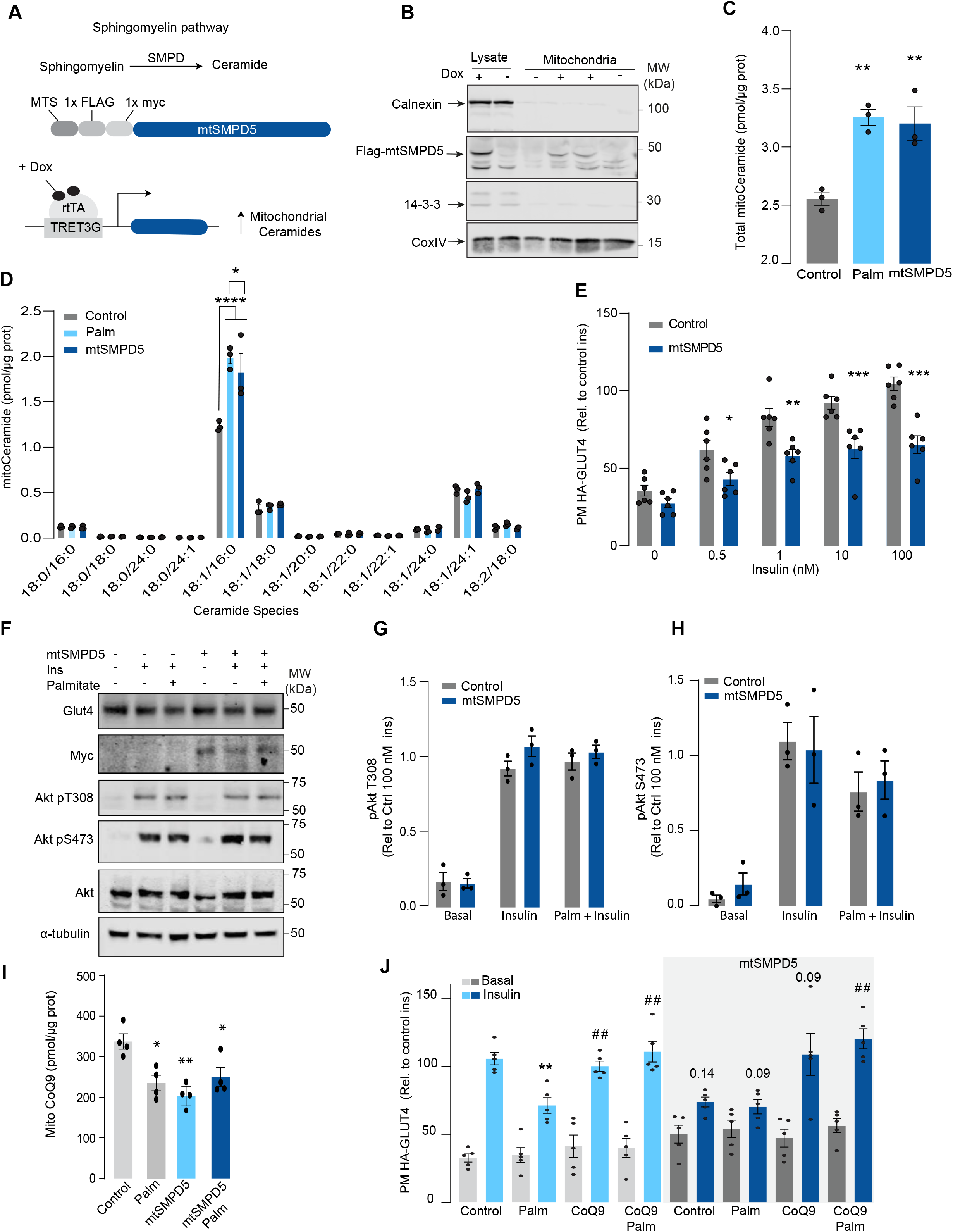
Mitochondrial overexpression of SMPD5 induces insulin resistance by lowering CoQ9 in L6-myotubes. A) Schematic representation of doxycycline-inducible overexpression of mitochondrial targeted sphingomyelinase 5 (SMPD5). L6-HA-GLUT4 myotubes were exposed to 1 μg/mL of Doxycycline from day 3 to day 6 of differentiation. Experiments were performed on day 7 of differentiation. B) Determination of SMPD5 expression in mitochondrial fraction obtained from L6-HA-GLUT4. Doxycycline was added where indicated. (C and D) Levels of endogenous ceramides in mitochondrial fraction from L6-HA-GLUT4 myotubes treated with BSA (Control) palmitate (150 μM for 16 h, palm) or doxycycline. Total (C) and specific (D) ceramide species were quantified. N = 3, mean ± S.E.M. *p< 0.05, ** p<0.01 and ****p< 0.0001 vs Control. E) Insulin-induced GLUT4 translocation in L6-HA-GLUT4 myotubes exposed to Doxycycline (1 μg/mL for 3 d). Plasma membrane GLUT4 (PM-GLUT4) abundance was normalised to 100 nM insulin-treated control cells. N = 6, mean ± S.E.M. *p< 0.05, **p<0.01, ***p<0.001 vs control ins. (F, G and H) L6-HA-GLUT4 myotubes were serum-starved after BSA (Control), Palmitate (150 μM for 16 h, palm) or Doxycycline (1 μg/mL for 3 d) treatment and acute insulin (Ins) was added where indicated. Phosphorylation status of indicated sites was assessed by immunoblot. Immunoblots were quantified by densitometry and normalised to insulin-treated control cells (indicated by dotted line). N = 3, mean ± S.E.M. * p<0.05, *** p<0.001 vs Basal I) CoQ9 level in mitochondrial fraction obtained from L6-HA-GLUT4 myotubes exposed to doxycycline (1 μg/mL for 3 d) N = 4, mean ± S.E.M. **p<0.001. J) Insulin-induced GLUT4 translocation in L6-HA-GLUT4 myotubes exposed to Doxycycline (1 μg/mL for 3 d). Control or doxycycline treated cells were exposed to BSA (control), Palmitate (150 μM for 16 h, palm) or CoQ9 (10 μM for 16 h). Plasma membrane GLUT4 (PM-GLUT4) abundance was normalised to insulin-treated control cells. N = 5, mean ± S.E.M. **p< 0.01 vs Control ins, ## p<0.01 vs palm ins

### Mitochondrial ceramides are necessary for palmitate-induced CoQ depletion and insulin resistance

Given increased mitochondrial ceramides are sufficient to induce CoQ depletion and insulin resistance, we next asked whether increased mitochondrial ceramides are necessary to drive these phenotypes. Using the doxycycline-Tet-On inducible system^40^ we overexpressed a mitochondrial-targeted Acid Ceramidase 1 (mtASAH1) in GLUT4-HA-L6 cells (GLUT4-HA-L6-mtASAH1) (Fig. 3A). ASAH1 degrades ceramides to fatty acid and sphingosine^44^. Hence, mitochondrial overexpression of ASAH1 was expected to selectively lower ceramides inside mitochondria. Doxycycline increased the abundance of mtASAH1 in the mitochondrial fraction, demonstrating the correct localisation of this construct (Fig. 3B). Furthermore, mtASAH1 induction prevented palmitate-induced mitochondrial ceramide accumulation (total levels and 18:1\16:0 ceramides) (Fig. 3C), indicating the enzyme was functioning as expected. Similar to our observations with mtSMD5 overexpression, mtASAH1 did not alter ceramide abundance in the whole cell lysate or mitochondrial sphingomyelin levels (Sup. Fig. 4 A&F). Notably, mtASAH1 overexpression protected cells from palmitate-induced insulin resistance without affecting basal insulin sensitivity (Fig. 3E). Similar results were observed using insulin-induced glycogen synthesis as an orthologous technique for GLUT4 translocation. These results provide additional evidence highlighting the role of dysfunctional mitochondria in muscle cell glucose metabolism (Sup. Fig. 5K). Importantly, mtASAH1 overexpression did not rescue insulin sensitivity in cells depleted of CoQ (2.5 mM 4NB for 24 h) supporting the notion that mitochondrial ceramides are upstream of CoQ (Fig. 3E). Neither palmitate nor mtASAH1 overexpression attenuated insulin-dependent Akt phosphorylation (Fig. 3 F-H) nor total GLUT4 abundance (Fig. 3F, Sup. Fig. 4B). Finally, mtASAH1 overexpression increased CoQ levels. In both control and mtASAH1 cells, palmitate induced a depletion of CoQ, however the levels in palmitate treated mtASAH1 cells remained similar to control untreated cells (Fig. 3I). This suggests that the absolute concentration of CoQ is crucial for insulin sensitivity, rather than the relative depletion compared to basal conditions, thus supporting the causal role of mitochondrial ceramide accumulation in reducing CoQ levels in insulin resistance.

**Figure 3.**
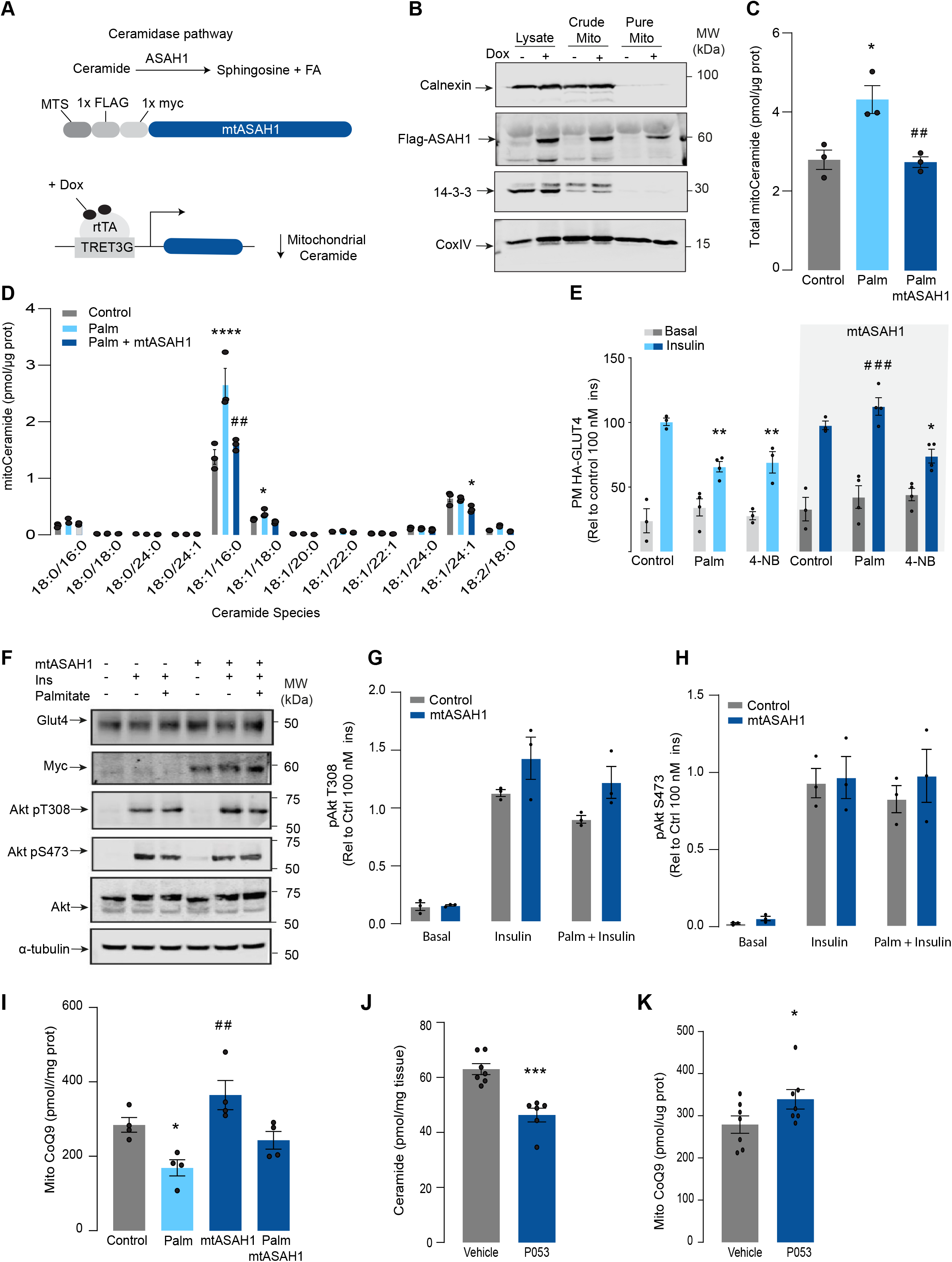
Mitochondrial overexpression of ASAH1 protects against insulin resistance and increases CoQ levels in L6-myotubes. A) Schematic representation of doxycycline-inducible overexpression of mitochondrial targeted Acid Ceramidase 1 (ASAH1). L6-HA-GLUT4 myotubes were exposed to 1 μg/mL of Doxycycline from day 3 to day 6 of differentiation. Experiments were performed on day 7 of differentiation. B) Determination of ASAH1 expression in mitochondrial fraction obtained from L6-HA-GLUT4. Doxycycline was added where indicated. (C and D) Endogenous ceramides levels in mitochondrial fraction from L6-HA-GLUT4 myotubes treated with BSA (Control) palmitate (150 μM for 16 h, palm) or doxycycline. Total (C) and specific (D) ceramide species were quantified. N = 3, mean ± S.E.M. *p< 0.05 vs Control, ## p<0.01 vs Palm E) Insulin-induced GLUT4 translocation in L6-HA-GLUT4 myotubes exposed to Doxycycline (1 μg/mL for 3 d). Control or doxycycline treated cells were exposed to BSA (control), Palmitate (150 μM for 16 h, palm) or 4NB (2.5 mM for 16 h). Plasma membrane GLUT4 (PM-GLUT4) abundance was normalised to insulin-treated control cells. N = 6, mean ± S.E.M. **p<0.01 vs Control ins, ### p<0.001 vs Palm ins (F, G and H) L6-HA-GLUT4 myotubes were serum-starved after BSA (Control), Palmitate (150 μM for 16 h, palm) or Doxycycline (1 μg/mL for 3 d) treatment and acute insulin (Ins) was added where indicated. Phosphorylation status of indicated sites was assessed by immunoblot. Immunoblots were quantified by densitometry and normalised to insulin-treated control cells (indicated by dotted line). N = 3, mean ± S.E.M. ***p<0.001 vs Basal, # p<0.05 vs Control ins. I) CoQ9 level in mitochondrial fraction obtained from L6-HA-GLUT4 myotubes exposed to doxycycline (1 μg/mL for 3 d). Control or doxycycline treated cells were exposed to BSA (control) or palmitate (150 μM for 16 h, palm) N = 4, mean ± S.E.M. *p<0.05 vs control, ## p<0.01 vs Palm. J) Levels of total ceramides in skeletal muscle of mice fed chow with vehicle or 5 mg/kg P053 for 6 wks. N = 7, Mean ± S.E.M. ***p< 0.001. K) Levels of CoQ in mitochondrial fraction isolated from skeletal muscle of mice fed chow with vehicle or 5 mg/kg P053 for 6 wks. N = 7, Mean ± S.E.M. *p<0.05.

In order to demonstrate the connection between ceramide and CoQ in vivo, we examined whether a reduction of ceramides in mouse skeletal muscle, using a Ceramide Synthase 1 (CerS1) inhibitor, would alter mitochondrial CoQ levels. Treatment of adult mice with the CerS1 inhibitor P053 for 6 wks selectively lowered muscle ceramides without affecting other lipid species (Fig 3J, Sup. Fig 5). Notably, CerS1 inhibition increased CoQ in mitochondrial fractions isolated from skeletal muscle (Fig. 3K), similar effect was observed in mice exposed to a high fat diet (HFD) for 5 wks (Supp. Fig. 4H-I). These animals exhibited an improvement in mitochondrial function and reduced muscle triglycerides and adiposity upon HFD (further phenotypic and metabolic characterization of these animals can be found in ^45^) demonstrating the existence of the ceramide/CoQ relationship in muscle in vivo.

### Mitochondrial ceramides induce depletion of the electron transport chain

We have established that both increased mitochondrial ceramides and a loss of mitochondrial CoQ are necessary for the induction of insulin resistance. As such these changes are likely to induce other mitochondrial defects. To gain insight into how increased mitochondrial ceramides drive changes in mitochondrial function we performed MS-based proteomics on L6 cells overexpressing mtSMPD5.

mtSMPD5-L6 myotubes were treated with doxycycline for various time points (2, 8, 24, 48 and 72 h) and positive induction was observed after 24 h of treatment (Fig. Sup. 6A). Subsequently, control, 24, and 72 h time points were selected for further studies. Mitochondria were purified via gradient separation and analysed using liquid chromatography-tandem mass spectrometry (LC-MS/MS) in data-independent acquisition (DIA) mode (Fig. 4A). Across control and mtSMPD5 cells we quantified 2501 proteins where 555 were annotated as mitochondrial proteins (MitoCarta 3.0 and uniprot localisation)^46^.

**Figure 4.**
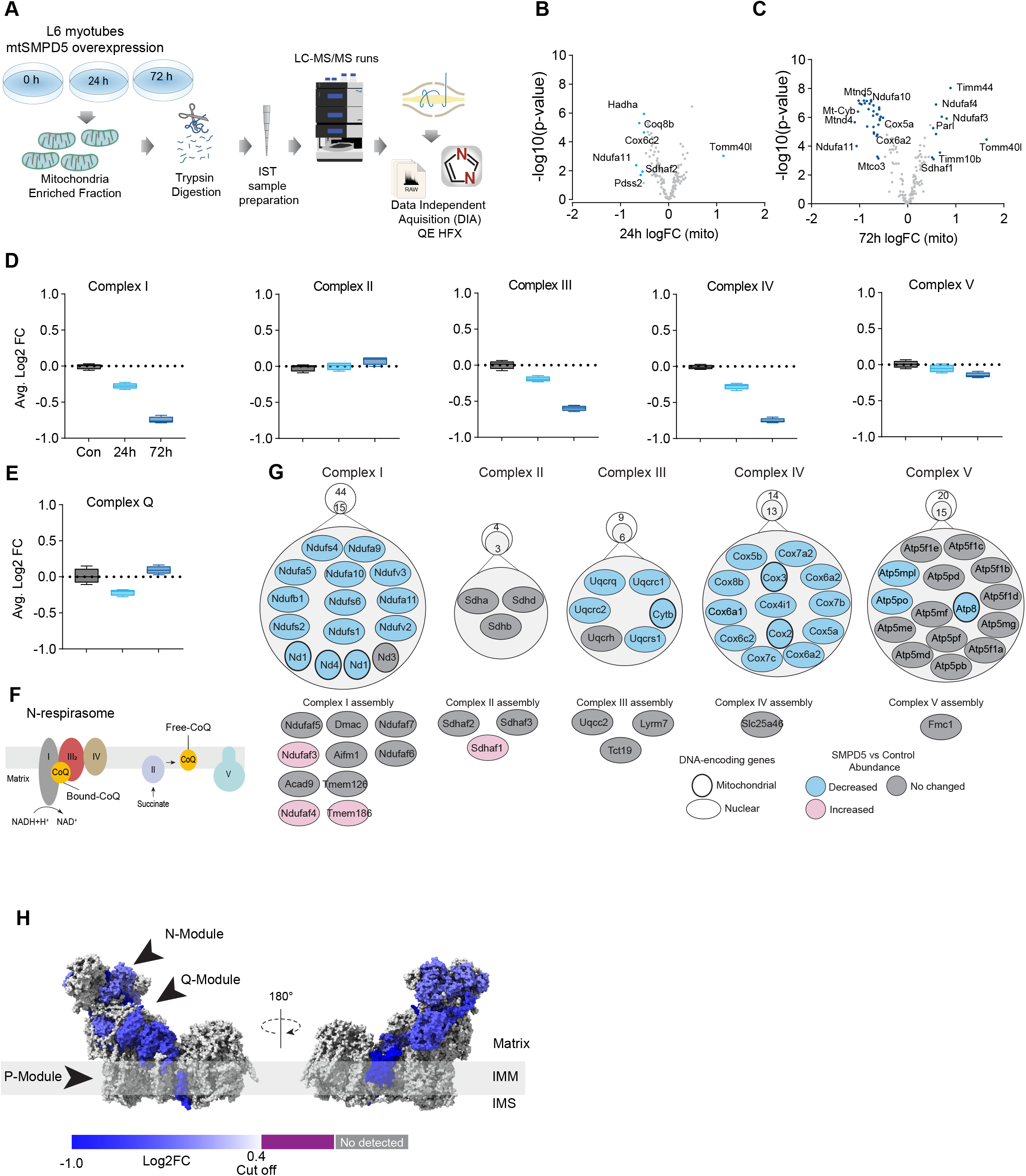
Mitochondrial ceramides induce a selective depletion of supercomplexes associated proteins in L6-myotubes. A) Workflow schematics. B & C) Pairwise comparisons of mitochondrial proteome between all four groups. Cut-off for -log10 adjusted p-value (-log10(p-value)) was set at 2 and Log2(FC) at 0.5 (blue). D) Quantification of the OXPHOS protein complexes generated by the summed abundance of all subunits within a given complex. E) Quantification of the Complex Q protein complexes generated by the summed abundance of all subunits within the complex. F) Schematics of CoQ distribution between CI-binding and free CoQ^82^ G) Schematics of OXPHOS subunits (top) and assembly factors (bottom) significatively up-regulated (light red), down-regulated (blue) and no change (grey) after 72 h of mtSMPD5 overexpression. H) Subunit levels for proteins after mtSMPD5 overexpression mapped to the complex I structure^48^. The colours were calculated with an in-house python script and the resultant model was rendered using ChimeraX. Grey, no detected; Purple, below cut off (Log2FC = 0.4).

Analysis of the proteome revealed that 9 and 19 % of mitochondrially annotated proteins were significantly changed at 24 and 72 h respectively (adj. p<0.05, Absolute log2 FC > 0.4) indicative of a temporal progression of changes following induction of mSMPD5 expression (Fig.4 B & C). 60 proteins were decreased at 72 h and of these 47% were functionally annotated as components of oxidative phosphorylation (OXPHOS (rank 1, 28/135 proteins). Within OXPHOS, we observed a significant depletion of the electron transport chain components (ETC). The ETC is composed of several complexes (complex I-V, CI-CV). In this dataset, CI (14/15 decreased), CIII (5/6 decreased) and CIV (13/13 decreased) but not CII (0/3 decreased) or CV (3/15 decreased) were depleted after mtSMPD5 overexpression (Fig. 4D). Despite the bulk downregulation of CI, III and IV, the assembly machinery associated with each complex was either upregulated or unchanged after mtSMPD5 overexpression (Fig. 4J) suggesting that mitochondrial ceramides somehow alter ETC stability. Intriguingly, neither CII nor CV were affected by mtSMPD5 suggesting that ceramides preferentially affect those ETC complexes that are part of structures known as supercomplexes (SCs)^47^. Importantly, as part of CoQ is found in SCs binding CI (Fig. 4F) we mapped the levels of individual subunits of CI onto the recently solved structure of bovine CI^48^. This revealed the loss of subunits around the N-module (Ndufs1, Ndufs4, Ndufs6, Ndufv2 & Ndufv3) and Q-module (Ndufa5 & Ndufs2) in CI (Fig. 4H). Importantly, CI downregulation was not associated with reduction in gene expression as shown in Sup. Fig. 6J. The N-module is essential for NADH oxidation, and the Q-module is where CoQ binds CI. Hence, loss of the Q-module might trigger a stoichiometric depletion of CoQ upon ceramide accumulation.

Of note, we observed a heterogeneous response of the mitochondrial proteome after mtSMPD5 overexpression. For instance, proteins associated with glucose oxidation and mitochondrial translation/transcription did not change after mtSMPD5 induction (Fig. Sup. 6D, F & G), proteins involved in fatty acid oxidation and OXPHOS were consistently downregulated after 24 h treatment (Fig. 4B, C, & D and Sup. Fig. 6E), proteins related with the mitochondrial import machinery were consistently upregulated (Fig. Sup. 6C) and proteins associated with CoQ production were transiently downregulated after 24 h induction (Fig. 4E).

### Mitochondrial ceramides impair mitochondrial function

Based on the ceramide-dependent depletion of ETC members, we hypothesised that mitochondrial ceramides would impair mitochondrial function. To test this, we evaluated several aspects of mitochondrial function upon mtSMPD5 overexpression. Mitochondrial respiration is broadly considered to be the best measure for describing mitochondrial activity^22,49^. Respiration was assessed in intact mtSMPD5-L6 myotubes treated with CoQ9 by Seahorse extracellular flux analysis. mtSMPD5 overexpression decreased basal and ATP-linked mitochondrial respiration (Fig. 5 A, B &C), as well as maximal, proton-leak and non-mitochondrial respiration (Fig. 5 A, D, E & F) suggesting that mitochondrial ceramides induce a generalised attenuation in mitochondrial function. Notably, we did not observe evidence of energy deficiency in our model (data not shown). Interestingly, CoQ9 supplementation partially recovered basal and ATP-linked mitochondrial respiration, suggesting that part of the mitochondrial defects are induced by CoQ9 depletion. The attenuation in mitochondrial respiration is consistent with a depletion of the ETC subunits observed in our proteomic dataset (Fig. 4). Since mitochondrial respiratory activity is limited by several factors including nutrient supply, bioenergetic demands, among others^50^ we tested whether the ETC generally or a specific respiratory complex was affected by mtSMPD5 overexpression. We measured the activity of the respiratory chain by providing substrates for each respiratory chain complex to permeabilized cells and analysed oxygen consumption. mtSMPD5 overexpressing cells exhibited attenuated mitochondrial respiration irrespective of the substrate provided (Fig. 5 G-L) supporting the notion that mitochondrial ceramides induce a generalised defect in mitochondrial respiration. In line with defective mitochondrial function, cells with mtSMPD5 overexpression also exhibited increased oxidative stress (Fig. 5M) measured by the redox sensitive dye MitoSOX. Interestingly, no difference in mitochondrial membrane potential was observed across conditions (Fig. 5N). Collectively, these data suggest that increased mitochondrial ceramides cause a loss of mitochondrial respiratory capacity and an increase in ROS production as a result of ETC depletion in L6-myotubes.

**Figure 5.**
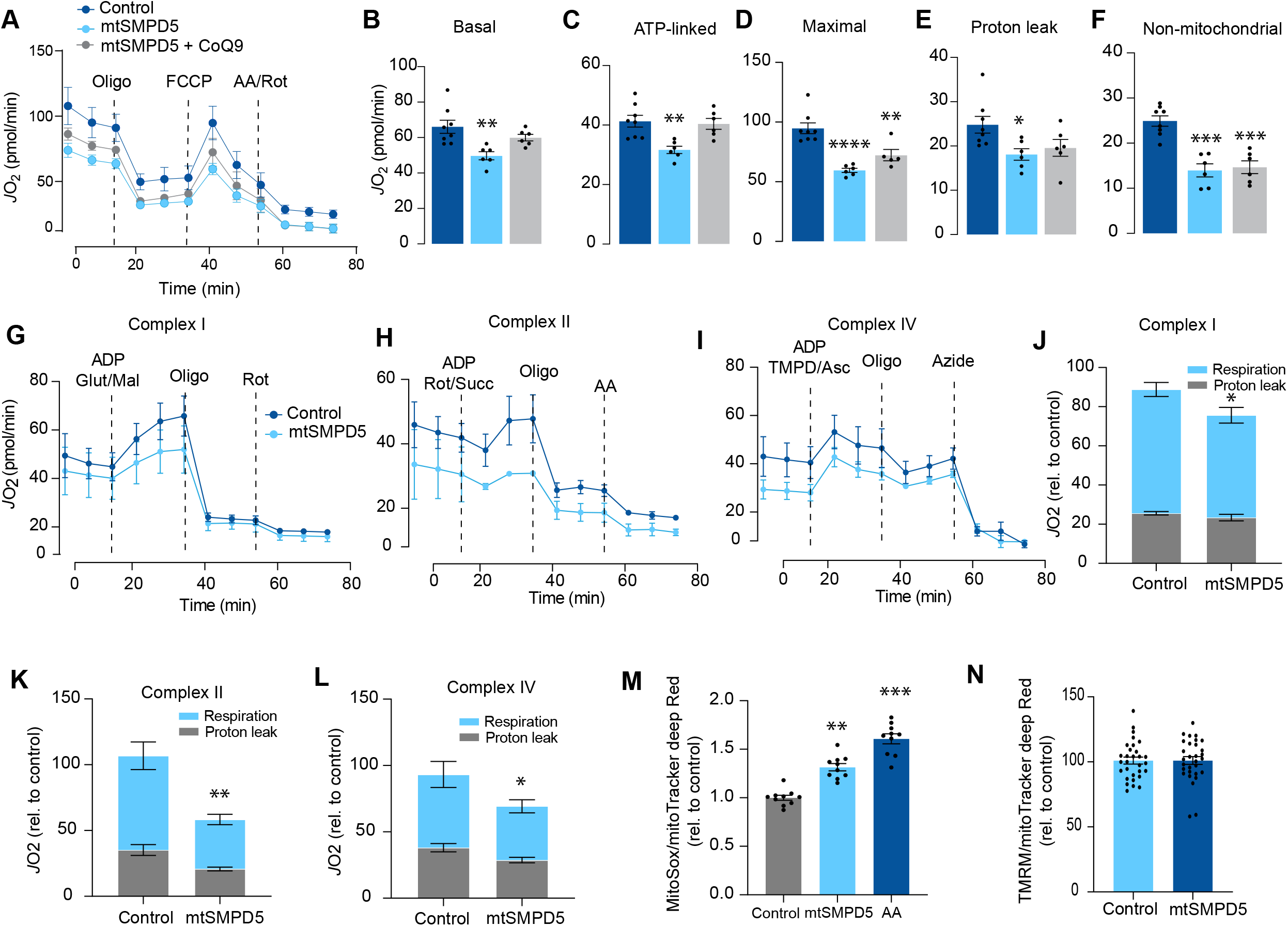
Mitochondrial ceramides impair mitochondrial function. A) mtSMPD5 overexpression decreases oxygen consumption rate (*J*O_2_) (means ± SEM; n = 6 - 8 biological replicates) measured by mitochondrial stress test. After 1 h no CO_2_ environment, cells were stimulated with oligomycin (Oligo), Carbonyl cyanide-p-trifluoromethoxyphenylhydrazon (FCCP) and antimycin A (AA) and Rotenote (Rot) at indicated time points. (means ± SEM; n = 6 - 8 biological replicates). B - F) Quantification of *J*O_2_ measured by mitochondrial stress test from Fig 6A as described in material and method. (means ± SEM; n = 6 - 8 biological replicates). * p<0.05, ** p<0.01, *** p<0.001 vs control G-L) mtSMPD5 overexpression diminishes respiratory CI (G & J), CII (H & K) and CIV (I & L). *J*O_2_ was performed in permeabilized cells supplemented with adenosine diphosphate (ADP) and CI to IV substrates (means ± SEM; n = 3 biological replicates). Mal, Malate; Glut, Glutamate; Rot, Rotenone; Succ, Succinate; TMPD, tetramethyl-phenylenediamine; Asc, Ascorbic acid; Oligo, Oligomycin. J, K and L are quantifications from graphs G, H and I respectively. * p<0.05, ** p<0.01 vs control. M) mtSMPD5 overexpression increased mitochondrial oxidative stress. Cells were loaded with the redox sensitive dye MitoSOX and the mitochondrial marker mitoTracker deep Red for 30 min before imaging in a confocal microscope (see method). (means ± SEM; n = 10 biological replicates). AA, Antimycin A. ** p<0.01, *** p<0.001 vs control N) mtSMPD5 overexpression does not alter mitochondrial membrane potential. Cells were loaded with the potentiometric dye Tetramethylrhodamine, Ethyl Ester, Perchlorate (TMRM^+^) in non quenching mode and the mitochondrial marker mitoTracker deep Red for 30 min before imaging in a confocal microscope (see method). (means ± SEM; n > 10 biological replicates).

### Association of mitochondrial proteome with insulin sensitivity and mitochondrial ceramides in human muscle

To further characterise the effect of mitochondrial ceramides on ETC abundance in a more physiological context we performed a cross-sectional study assessing the mitochondrial lipid profile and protein abundance in muscle biopsies obtained from four groups of people (athletes, lean, obese and type 2 diabetics (T2D). The demographic information and detailed lipidomic analysis of these individuals was previously reported^30^, Fig. Sup. 7 A). In line with our *in-vitro* data, long tail ceramides (C18:0) in the mitochondria/ Endoplasmic reticulum (ER) enriched fraction, but not whole tissue, were inversely correlated with muscle insulin sensitivity^30^. To expand this observation, we employed proteomics analysis of the mitochondrial/ER fraction from the same subjects (Fig. 6A). A total of 2,058 unique protein groups were quantified in at least one sample, where 571 were annotated as mitochondrial associated proteins (Human MitoCarta 3.0)^46^. After filtering (proteins in >50% of samples within each group), 492 mitochondrial proteins were reliably quantified across 67 samples (Fig. 6B). We noted that the mitochondrial fraction from athletes were enriched for mitochondrial proteins, and this could be corrected by global median normalisation (Fig. Sup. 7D). Pairwise comparison of the mitochondrial proteome between all groups revealed differences between groups, although relatively small in effect size (Fig. 6C & D). For instance, 16% of all mitochondrial proteins were significantly different between T2D and athletes (Fig. 6C), however 56% (45/80) of these proteins were changed by less than 1.5-fold. This trend was even stronger when comparing the obese group to the athletes, where 18% of mitochondrial proteins were changed, but ∼80% were changed less than 1.5-fold. Gene set enrichment revealed a highly significant general trend following the difference in insulin sensitivity (measure by the rate of glucose disappearance -Rd- using a stable isotope - [6,6-^2^H_2_]glucose - during a hyperinsulinemic-euglycemic clamp ^30^, where TCA cycle and respiratory electron transport and Complex I biogenesis were enriched as follows: Athletes > Lean > Obese > T2D (Sup. Table 2). Of note, the T2D group had an enrichment of mitochondrial translation compared to the obese group.

**Figure 6.**
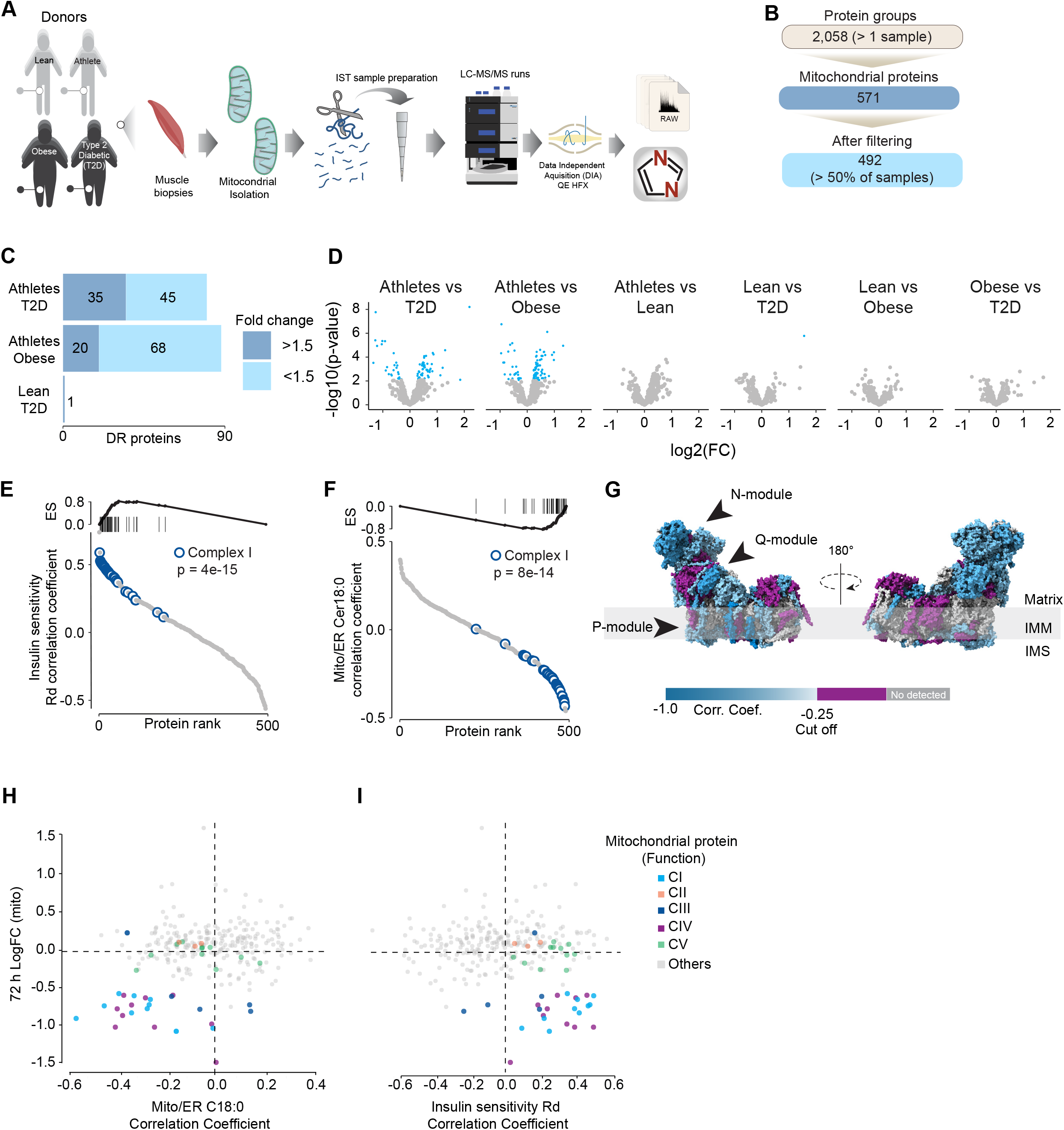
Mitochondrial proteome profiling associates complex I with muscle insulin sensitivity. A) Quantification of proteins across samples. B) Number of significant deferential regulated (DR) proteins by pairwise comparison. Groups not shown have no significantly regulated proteins after correcting for multiple testing. C) Gene set enrichment between all comparisons. D) Relative protein abundance in isolated mitochondria from human skeletal muscle muscle cells. Comparisons are shown on top of each graph. Light blue, significatively regulated proteins (-log10(p-val) = 2). E) Proteins rank against rate glucose disappearance during clamp (Rd) correlation. Proteins within complex I of the electron transport chain are highly significant with Rd. F) Proteins rank against mitochondrial ceramide (Cer) 18:0 abundance. Proteins within complex I of the electron transport chain are highly significant with Cer18:0. G) Subunit levels associated with mitochondrial ceramides mapped to the complex I structure ^48^. The colours were calculated with an in-house python script and the resultant model was rendered using ChimeraX. Grey, no detected; Purple, below cut off (Corr. Coef. = - 0.25 to 0.25). H) Quantification of the mitochondrial proteins generated by the summed abundance of all subunits associated with a specific function from mtSMPD5-L6 myotubes after 72 h vs summed mitochondrial proteins (function) associated with either C18:0 ceramide (left panel) or insulin sensitivity (right panel) from human samples.

To further explore the relationship between mitochondria and insulin sensitivity, the mitochondrial proteome was correlated to the muscle insulin sensitivity measured using ^2^H_2_ glucose Rd. As a group, all detected proteins within CI of the ETC were highly correlated with muscle insulin sensitivity (p = 4e-15) (Fig. 6E), and to a lesser extent proteins within CIV (p=0.08) and CV (p= 0.09; Fig. Sup. 7E). The abundance of CII and CIII, together with the small and large mitochondrial ribosome subunits, were not associated with insulin sensitivity across all the samples (Fig. Sup. 7F). Next, we determined the association between the mitochondrial proteome and the levels of C18:0 ceramide in the mitochondria/ER fraction. In line with our previous observations, as a group, CI proteins were inversely correlated with mitochondrial ceramides (p = 8e-14) and no association was observed between CII and C18:0 ceramides across samples (Fig. 6F). Furthermore, components of CIV were also negatively correlated with mitochondrial ceramides although to a lesser extent (Sup. Fig. 7F, p = 0.026) and CV was not associated with mitochondrial ceramides (Sup. Fig. 7F, p = 0.737). According to these results, ETC subunits exhibit differential sensitivity to mitochondrial ceramides, with CI subunits being the most sensitive in human muscle. To uncover structural changes in CI that could correlate with increased ceramide we mapped those proteins significatively associated with mitochondrial ceramides to the bovine CI structure^48^. Consistent with L6-mtSMPD5 myotubes, the N and Q modules were the regions with the most negative associated subunits with mitochondria ceramides in human muscle (Fig. 6G). To determine the conservation in the changes in the mitochondrial proteome induced by increased ceramides, we compared the proteomes of mtSMPD5-L6-myotubes (72 h after induction) and human muscle biopsies. We observed that across the two datasets, CI and CIV subunits were downregulated after mtSMPD5 overexpression and were negatively associated with C18 ceramides in human samples (Fig. 6H). In turn, CI and CIV were positively associated with muscle insulin sensitivity (Fig. 6I), suggesting that these ETC subunits exhibited a conserved sensitivity to ceramide accumulation with a potential role in insulin sensitivity.

## Discussion

Insulin resistance is characterised by attenuated insulin-dependent glucose uptake in relevant target tissues, such as muscle and fat, and it plays a central role in cardiometabolic diseases^3^. In skeletal muscle, mitochondrial ceramides have been linked to insulin resistance^30^, however, to date no direct link connecting mitochondrial ceramides with insulin resistance has been established. Furthermore, CoQ depletion and defective mitochondria have also been independently associated with insulin resistance^6,7^. In the current study we present evidence suggesting that these factors are mechanistically linked inside mitochondria. Our data demonstrate that increased mitochondrial ceramides are both necessary and sufficient to induce insulin resistance in skeletal muscle. This is likely a function of increased ROS production that results from the specific depletion of the OXPHOS subunits and the concomitant loss of CoQ. Analysis of the human muscle mitochondrial proteome strongly supports mitochondrial ceramide linked changes in the OXPHOS machinery as major drivers of insulin sensitivity. In this study, we mainly utilised L6-myotubes, which share many important characteristics with primary muscle fibres. Both types of cells exhibit high sensitivity to insulin and respond similarly to maximal doses of insulin, with GLUT4 translocation stimulated between 2 to 4 times over basal levels in response to 100 nM insulin (as shown in Fig. 1-4,^51,52^). Additionally, mitochondrial respiration in L6-myotubes have a similar sensitivity to mitochondrial poisons, as observed in primary muscle fibres (as shown in Fig. 5,^53^). Finally, inhibiting ceramide production increases CoQ levels in both L6-myotubes and adult muscle tissue (as shown in Fig. 2-3). Therefore, L6-myotubes possess the necessary metabolic features to investigate the role of mitochondria in insulin resistance, and this relationship is likely applicable to primary muscle fibres.

Many stressors, including chronic inflammation and anticancer drugs, stimulate endogenous ceramide generation^54^ and CoQ depletion in mitochondria^55,56^. Nevertheless, experimental evidence testing the link between these molecules has been lacking. We observed that increased mitochondrial ceramides drive a depletion of mitochondrial CoQ leading to insulin resistance (Fig. 1 & 2), and that reducing mitochondrial ceramide protects against the loss of CoQ and IR (Fig. 3). These findings align with our earlier observations demonstrating that mice exposed to HFHSD exhibit mitochondrial CoQ depletion in skeletal muscle ^6^. Given that CoQ supplementation is sufficient to overcome ceramide induced-IR (Fig. 1 & 2), but a reduction of mitochondrial ceramide does not overcome a loss of CoQ (Fig. 3), our data support a pathway whereby an increase in mitochondrial ceramides precedes loss of CoQ. Interestingly, inhibition of CoQ synthesis also increased ceramides, suggesting a bidirectionality to the ceramide-CoQ nexus. That said, this effect was modest (Fig. 1) and we cannot exclude off target effects of the inhibitor. It is possible that CoQ directly controls ceramide turnover^34^ or alternatively that CoQ inside mitochondria is necessary for fatty acid oxidation^12^ and CoQ depletion triggers lipid overload in the cytoplasm promoting ceramide production^35^. Further studies will be needed to determine how CoQ depletion promotes ceramide accumulation.

Our proteomics analysis revealed that the loss of CoQ parallels a loss of mitochondrial ETC complexes CI, CIII and CIV. These are known to form supercomplexes or respirasomes where ∼25 - 35 % of CoQ is localised in mammals ^57,16^. This bulk downregulation of the respirasome induced by ceramides may lead to CoQ depletion. The observation that both palmitate and SMPD5 overexpression trigger CoQ depletion without additive effects support the notion that ceramides may trigger the depletion of a specific CoQ9 pool localised within the inner mitochondrial membrane. Despite the significant impact of ceramide on mitochondrial respiration, we did not observe any indications of cell damage in any of the treatments, suggesting that our models are not explained by toxicity and increased cell death (Sup. Fig. 2H & J). Whilst the physiological role of respirasomes is still a subject of discussion, it has been suggested that they may enhance energy generation by optimising electron flow while reducing production of ROS^58,59^ and therefore their loss can be predicted to increase ROS generation. The 2 major mechanisms that might account for the loss of the respirasomes are decreased synthesis or increased degradation. Proteomics data suggests no deficiency in the OXPHOS biosynthetic machinery or assembly proteins and an increase in the machinery for protein import (Fig. 4). In addition, the absence of mRNA downregulation in mtSMPD5 overexpressing cells strongly suggests that at least a portion of the observed protein depletion within CI is attributed to diminished protein stability. It therefore seems reasonable to speculate that the loss of these mitochondrial complexes is driven by increased degradation. Interestingly, pharmacological CIII inhibition leads to respirasome degradation via oxidative stress produced by reverse electron transfer (RET)^60^. Since ceramides can directly inhibit CIII^61^ it is possible that a similar mechanism mediates the effect of ceramides on the respirasome (Fig. 5). This suggests that defective respirasome activity (e.g. induced by ceramides) triggers ROS, which over time depletes respirasome subunits and a stoichiometric CoQ depletion, leading to further ROS production as a consequence. Another possibility is that, because of its highly hydrophobic nature, ceramides impact membrane fluidity promoting a gel/fluid phase transition^62^. These alterations in membrane fluidity could decrease respirasome stability. It is likely that bound lipids stabilise the interactions between the complexes in the respirasome and that this is impaired by ceramides. In fact, bound lipid molecules are observed in the structure of the porcine respirasome^63^ and the isolated bovine CI^63^ but none of the lipids identified thus far directly bridge different complexes. In order to understand the role of lipids in stabilising respirasomes and the role of ceramides in such stabilisation, higher-resolution structures will be required^64^.

The current studies pose a number of key unanswered questions. First, how does ceramide accumulate in mitochondria in insulin resistance? This could involve transfer from a different subcellular compartment^65,66^ or *in situ* mitochondrial ceramide synthesis. Consistent with the latter, previous studies have suggested that various enzymes involved in ceramide metabolism are specifically found in mitochondria^41,67–71^. Notably, CerS1-derived ceramide induces insulin-resistance in skeletal muscle^72^. Although this enzyme has not been reported as a mitochondrial protein, it can be transferred from the endoplasmic reticulum surface to the mitochondria under cellular stress in metabolically active tissues such as muscle and brain^73^. This provides a potential mechanism where cellular stress, like nutrient overload, may induce transfer of CerS1 to mitochondria, increasing mitochondrial ceramide to trigger insulin resistance.

A further question is how ceramide regulates insulin sensitivity. We present evidence that mitochondrial dysfunction precedes insulin resistance. However, previous studies have failed to observe changes in mitochondrial morphology, respiration or ETC components during early stages of insulin resistance^22,74^. However, in many cases such studies fail to document changes in insulin-dependent glucose metabolism in the same tissue as was used for assessment of mitochondrial function. This is crucial because we and others do not observe impaired insulin action in all muscles from high fat fed mice for example ^6,9^. In addition, surrogate measures such as insulin-stimulated Akt phosphorylation may not accurately reflect tissue specific insulin action as demonstrated in figure 1C. Thus, further work is required to clarify some of these inconsistencies. We observed that mitochondrial ceramides were associated with the loss of CoQ, increased production of mitochondrial ROS and impaired mitochondrial respiration^6,9,10^. As we discussed above, this is likely a direct result of respirasome depletion. The molecular linkage between ROS production and IR remains unknown. Early studies suggested that ceramides and ROS impaired canonical insulin signalling^24–26^, however, our current data do not support this, with the caveat that these were static signalling measures. One possibility is the release of a signalling molecule from the mitochondria that impairs insulin action^75^. The mitochondrial permeability transition pore (mPTP) is an attractive candidate for this release since its activity is increased by mitochondrial ROS^76^ and ceramides^77^. It has been shown that mPTP inhibition protects against insulin resistance in either palmitate- or ceramide-induced L6-myotubes and mice on a high-fat diet^78^. Furthermore, mPTP deletion in the liver protects against liver steatosis and insulin resistance in mice^79^. Strikingly, CoQ is an antioxidant and also an inhibitor of mPTP suggesting that part of the protective mechanism of CoQ may involve the mPTP^80^. Because CoQ can accumulate in various intracellular compartments, it’s important to consider that its impact on insulin resistance might be due to its overall antioxidant properties rather than being limited to a mitochondrial effect. Excitingly, mtSMPD5 increased the abundance of mPTP associated proteins suggesting a role of this pore in ceramide induced insulin resistance (Sup. Fig. 6E). It is also possible that ceramides generated within mitochondria in SMPD5 cells leak out from the mitochondria into other membranes (e.g. PM and GLUT4vesicles) affecting other aspects of GLUT4 trafficking and insulin action. However, the observation that ASAH1 overexpression reversed IR without affecting whole cell ceramides argues against this possibility.

Ultimately, the significant challenge for the field is the discovery of the unknown factor(s) released from mitochondria that cause insulin resistance, their molecular target(s), and the transduction mechanism(s).

The observations described above led us to speculate on whether there is a teleological reason for why these mitochondrial perturbations occur and why they drive insulin resistance? Under conditions of stress, nutrient incorporation into the cell needs to be adjusted to keep the balance between energy supply and utilisation. In situations where the mitochondrial respirasome is depleted, the mitochondria’s ability to oxidise nutrients can be easily overwhelmed without a corresponding reduction in nutrient uptake. In this scenario, insulin resistance may be a protective mechanism to prevent mitochondrial nutrient oversupply^9^. Beyond nutrient uptake, the respirasome depletion could also affect the ability of the mitochondria to switch between different energy substrates depending on fuel availability, named “metabolic Inflexibility”^81^ this mechanism may potentially play a role in the ectopic lipid accumulation seen in individuals with obesity, a condition linked with cardio-metabolic disease.

In summary, our results provide evidence for the existence of a mechanism inside mitochondria connecting ceramides, mitochondrial respiratory complexes, CoQ and mitochondrial dysfunction as part of a core pathway leading to insulin resistance. We identified that CoQ depletion links ceramides with insulin resistance and define the respirasome as a critical connection between ceramides and mitochondrial dysfunction. While many pieces of the puzzle remain to be solved, identifying the temporal link between ceramide, mitochondrial dysfunction and CoQ in mitochondria is an important step forward in understanding insulin resistance and other human diseases affecting mitochondrial function.

## Acknowledgments

This work was supported by National Health and Medical Research Council (NHMRC) Project Grants: GNT1120201 and GNT1061122 to D.E.J.; GNT2013621 to J.G.B, D.E.J. and A.D.V; and GNT112613 to N.T., J.M. and A.D. B.C.B received a research grant from Eli Lilly and from NIH R01DK111559. D.E.J. is an Australian Research Council (ARC) Laureate Fellow. J.G.B and A.D.V. were supported by the Diabetes Australian Research Program (Y22G-DIAA) and the Mitochondrial Foundation (Mitofoundation, G057). The content is solely the responsibility of the authors and does not necessarily represent the official views of the NHMRC or ARC. The authors also acknowledge the facilities, and the scientific and technical assistance of Sydney Cytometry and the Sydney Mass Spectrometry Facility, at the Charles Perkins Centre, University of Sydney. The authors are extremely grateful to Scott A. Summers, William L. Holland and Navdeep S Chandel for their thoughtful discussion of the project.

## Author contributions

Conceptualization; A.D.V., J.G.B., B.C.B, J.T.B., and D.E.J. Formal analysis; A.D.V., J.G.B., A.D., J.K. Investigation; A.D.V., K.C.C., J.G.B., J.K. Methodology; A.D.V., J.G.B., K.C.C., L.C., J.K., N.T., X.Y.L., J.M., A.D., A.G., S.Z., K.A.Z.B., A.R., B.C.B., S.M., M.A.A., and J.T.B. Writing – original draft; A.D.V, J.G.B. and D.E.J. Writing - review & editing; all authors.

## Supplementary Figures

**Figure S1.**
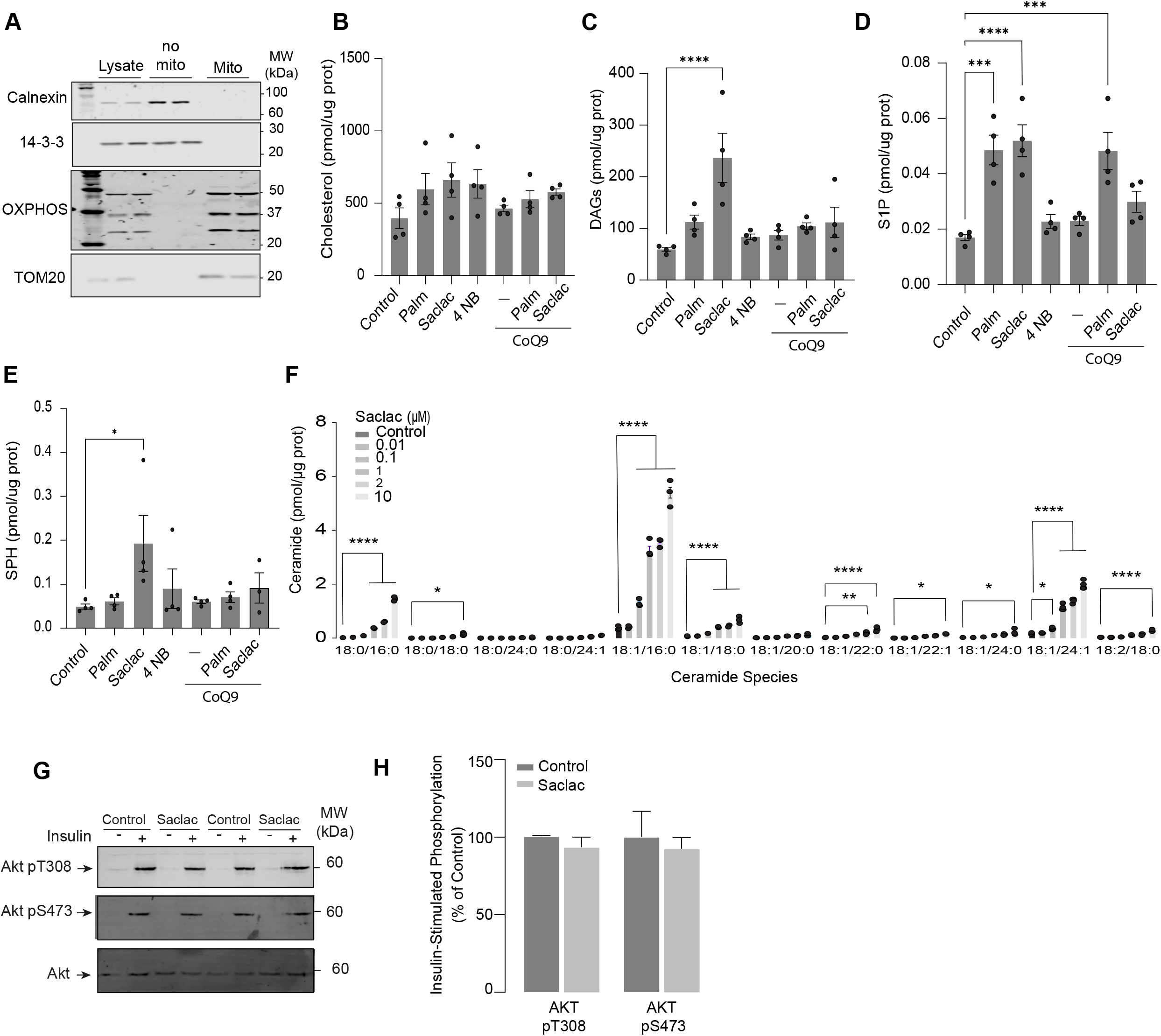
A) Mitochondrial enriched fraction from L6-myotubes. Calnexin was used as a marker of endoplasmic reticulum. OXPHOS: oxidative phosphorylation system. Cholesterol (B), Diacylglycerol (DAGs) (C), Sphingosine-1 phosphate (S1P) (D), and Sphingosine (SPH) (E) abundance in L6 myotubes exposed to different compounds (as indicated). Lipid abundance was determined by lipidomics and normalised against protein concentration. N = 4, Mean ± S.E.M. ***p< 0.001, ****p< 0.0001. F) Concentration of endogenous ceramide in L6-HA-GLUT4 myotubes treated for 24 h with EtOH (control) or different concentrations of Saclac. N = 3, mean ± S.E.M. *p<0.05. **p< 0.01, ****p< 0.0001. G-H) L6-HA-GLUT4 myotubes were serum-starved after EtOH (Control) or Saclac (10 μM) treatment (for 24 h) and acute insulin (Ins) was added where indicated. Phosphorylation status of indicated sites was assessed by immunoblot. Immunoblots were quantified by densitometry and normalised to insulin-treated control cells. N = 2, mean ± S.E.M.

**Figure S2.**
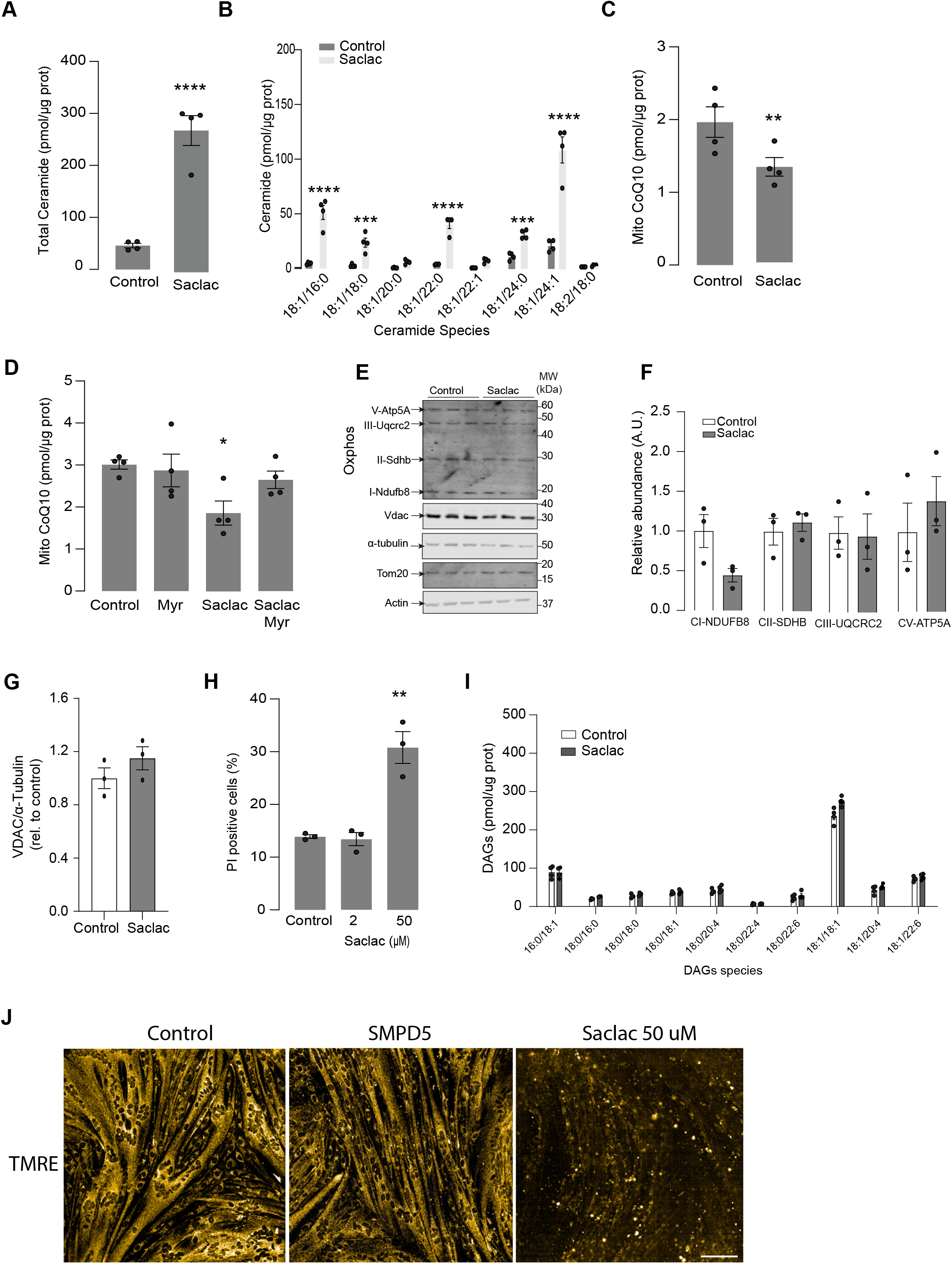
(A and B) Total (A) and specific (B) ceramide species quantified in HeLa cells treated for 24 h with Saclac (2uM for 24 h) or vehicle control (EtOH, Control) as indicated. N = 4, mean ± S.E.M. **p< 0.01, ****p< 0.0001 vs control C) CoQ10 levels in the mitochondrial fraction obtained from HeLa cells exposed to different concentrations of Saclac or vehicle control. N = 4, mean ± S.E.M. *p<0.05 D) CoQ10 levels in the mitochondrial fraction obtained from HeLa cells exposed to different concentrations of Saclac (2 μM for 24 h) or control in presence or absence of myriocin (10 μM for 16 h). N = 4, mean ± S.E.M. *p<0.05. E-G) Mitochondrial abundance markers determined by Western Blot in HeLA cells exposed to 2 uM of Saclac for 24 h. F - G) Immunoblots were quantified by densitometry and normalised to control cells. N = 3, Mean ± S.E.M H) Percentage of non-viable HeLa cells determined by propidium iodide (PI) staining and microscopy, following a 24 h treatment with Saclac. N = 3, mean ± S.E.M. **p<0.01. I) specific diacylglycerol species were quantified in HeLA cells exposed to Saclac vs control. Lipid abundance was determined by lipidomics and normalised against protein concentration. N = 4, Mean ± S.E.M. *p< 0.001, ****p< 0.0001. J) L6 myotubes were loaded with TMRE (20 nM) for 20 min in control cells, SMPD5 overexpressing cells and cells treated with Saclac (50 uM) for 24 h.

**Figure S3.**
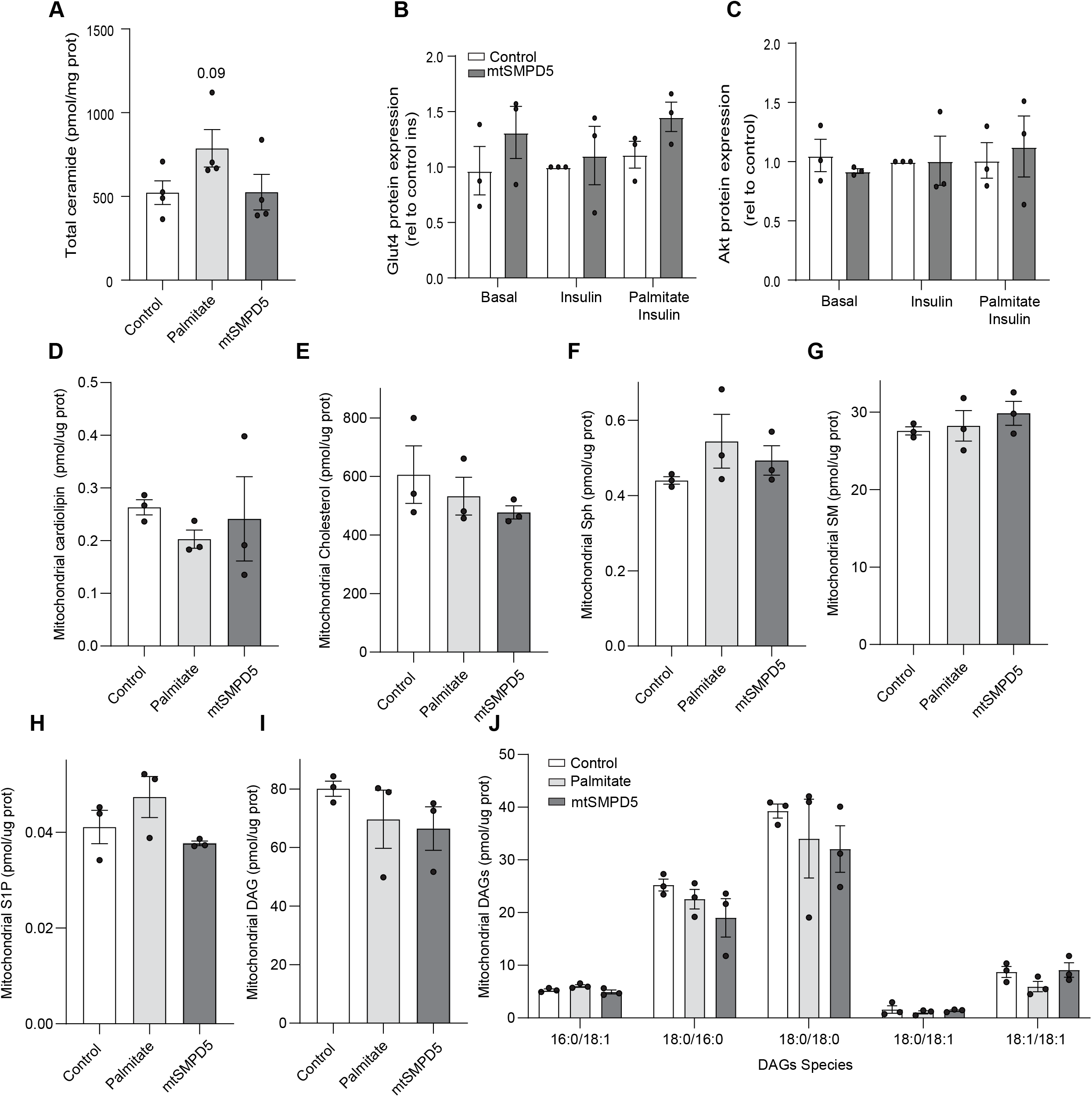
A) Total ceramide levels extracted from whole lysate of L6-myotubes exposed to different treatments (as indicated). B-C) Densitometric analysis of Figure 3F of selected proteins (As indicated). Mitochondrial levels of cardiolipin (D), cholesterol (E), Sphingosine (F), sphingomyelin (G), Sphingosine-1 phosphate (H) and diacylglycerol (I - J) (as indicated) were determined by lipidomics and normalised against mitochondrial protein concentration. Cells were exposed to doxycycline for 3 d for SMPD5 induction. N= 3, Mean ± S.E.M. ****p< 0.0001.

**Figure S4.**
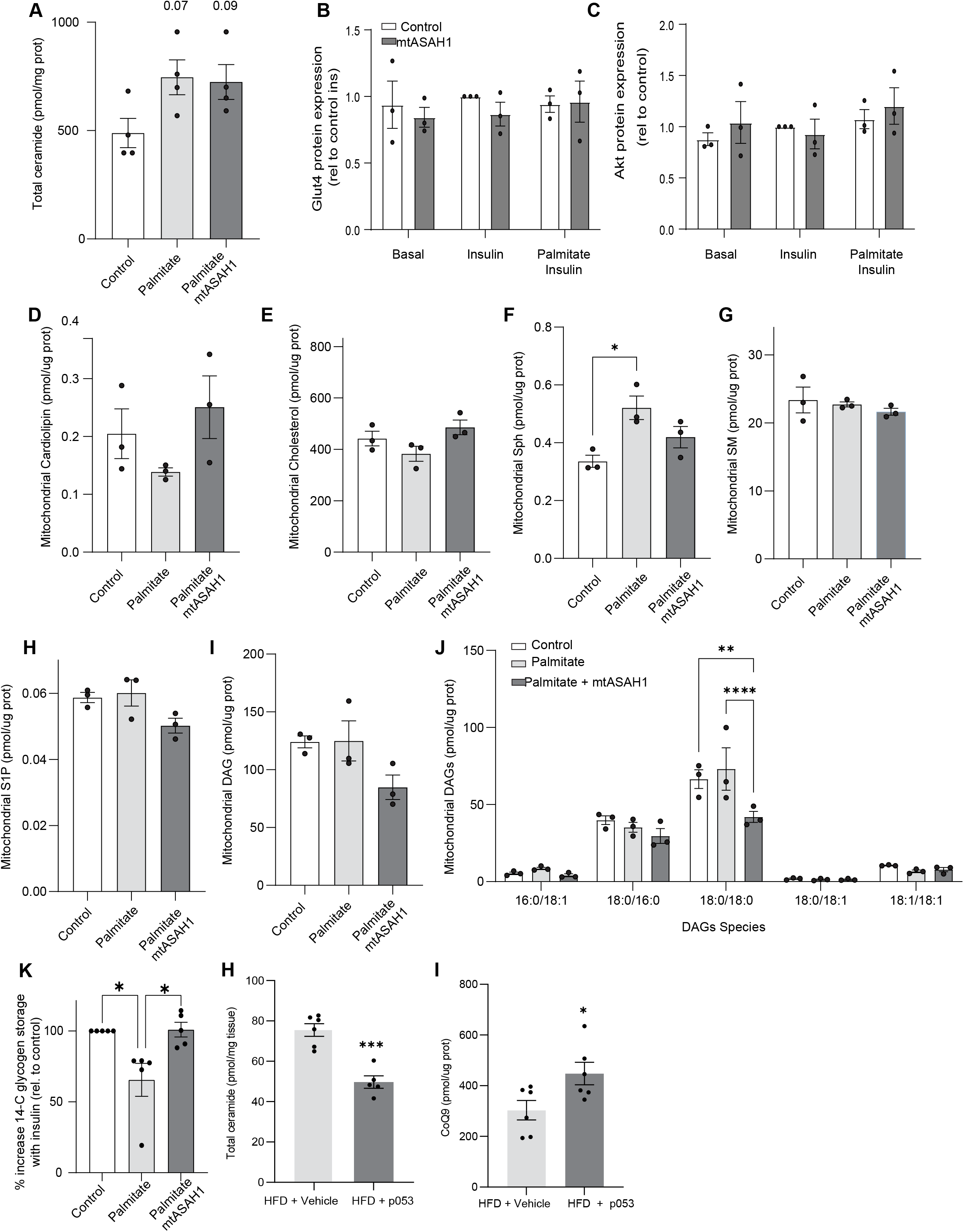
A) Total ceramide levels extracted from whole lysate of L6-myotubes exposed to different treatments (as indicated). B-C) Densitometric analysis of Figure 3F of selected proteins (As indicated). Mitochondrial levels of cardiolipin (D), cholesterol (E), Sphingosine (F), sphingomyelin (G), Sphingosine-1 phosphate (H) and diacylglycerol (I - J) (as indicated) were determined by lipidomics and normalised against mitochondrial protein concentration. Cells were exposed to doxycycline for 3 d for ASAH1 induction. N= 3, Mean ± S.E.M. *p<0.05, **p<0.01, ****p< 0.0001. K) Insulin-induced glycogen synthesis. L6-myotubes were exposed to different treatments (as indicated) and glycogen storage was measured by 14-C glucose (as indicated in material and method section). Data is expressed as a percentage of increased 14-glycogen after insulin stimulation relative to control cells. N = 4, mean ± S.E.M. *p<0.05. H-I) Total ceramides (H) and mitochondrial CoQ levels (I) in skeletal muscle from mice exposed to a high-fat diet and exposed to either vehicle (DMSO) or the inhibitor of CerS1 P053 (5 mg/Kg) in drinking water for 6 wks as described in ^45^. N = 6, mean ± S.E.M. *p<0.05, *** p<0.001 vs control.

**Figure S5.**
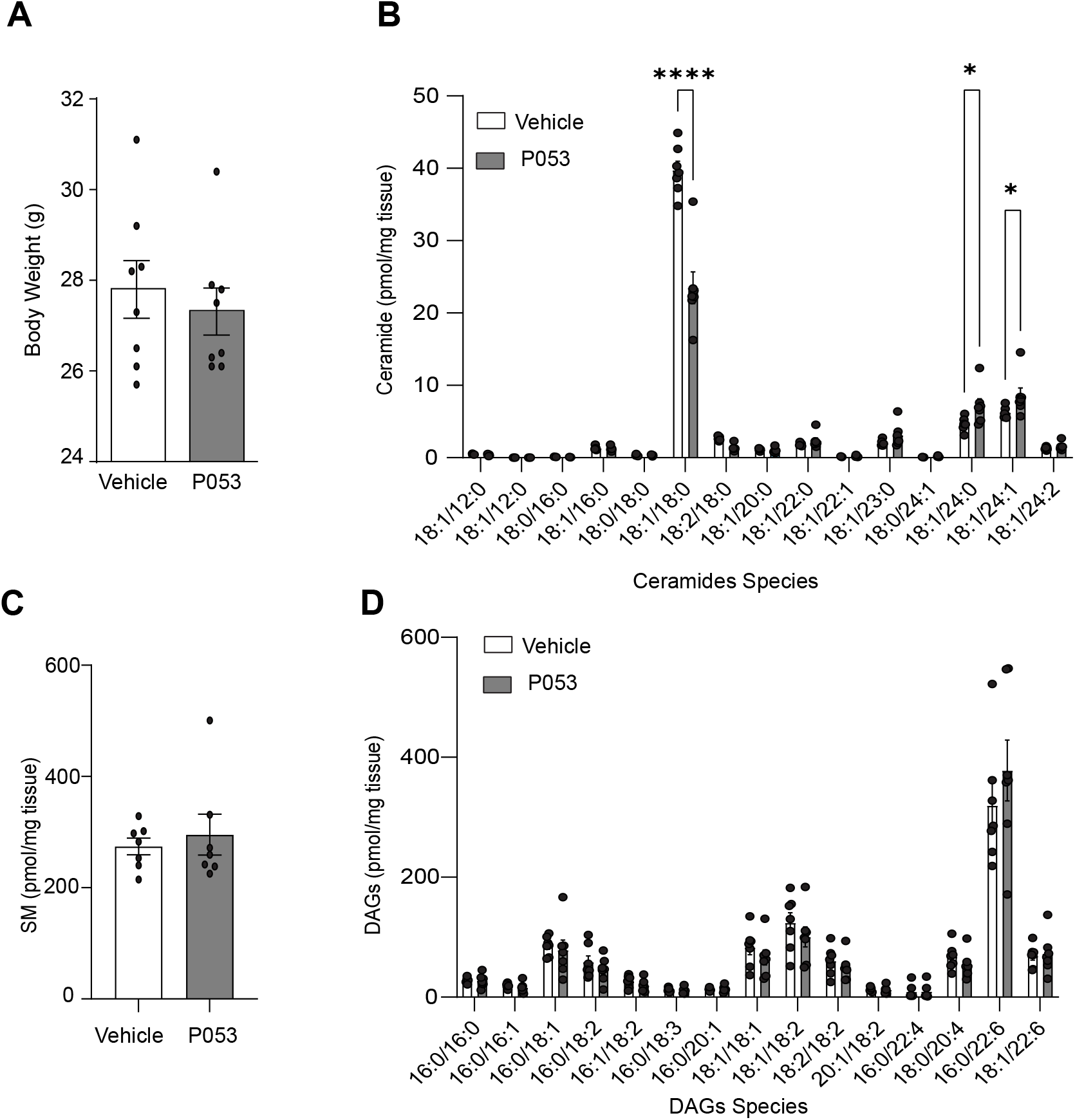
A) Body weight of mice exposed to either vehicle (DMSO) or the inhibitor of CerS1 P053 (5 mg/Kg) in drinking water for 6 wk. B) Ceramide, C) Sphingomyelin and D) Diacylglycerol species were determined in skeletal muscle by lipidomics and normalised against mg of tissue. N= 6, Mean ± S.E.M. *p<0.05, ****p< 0.0001.

**Figure S6.**
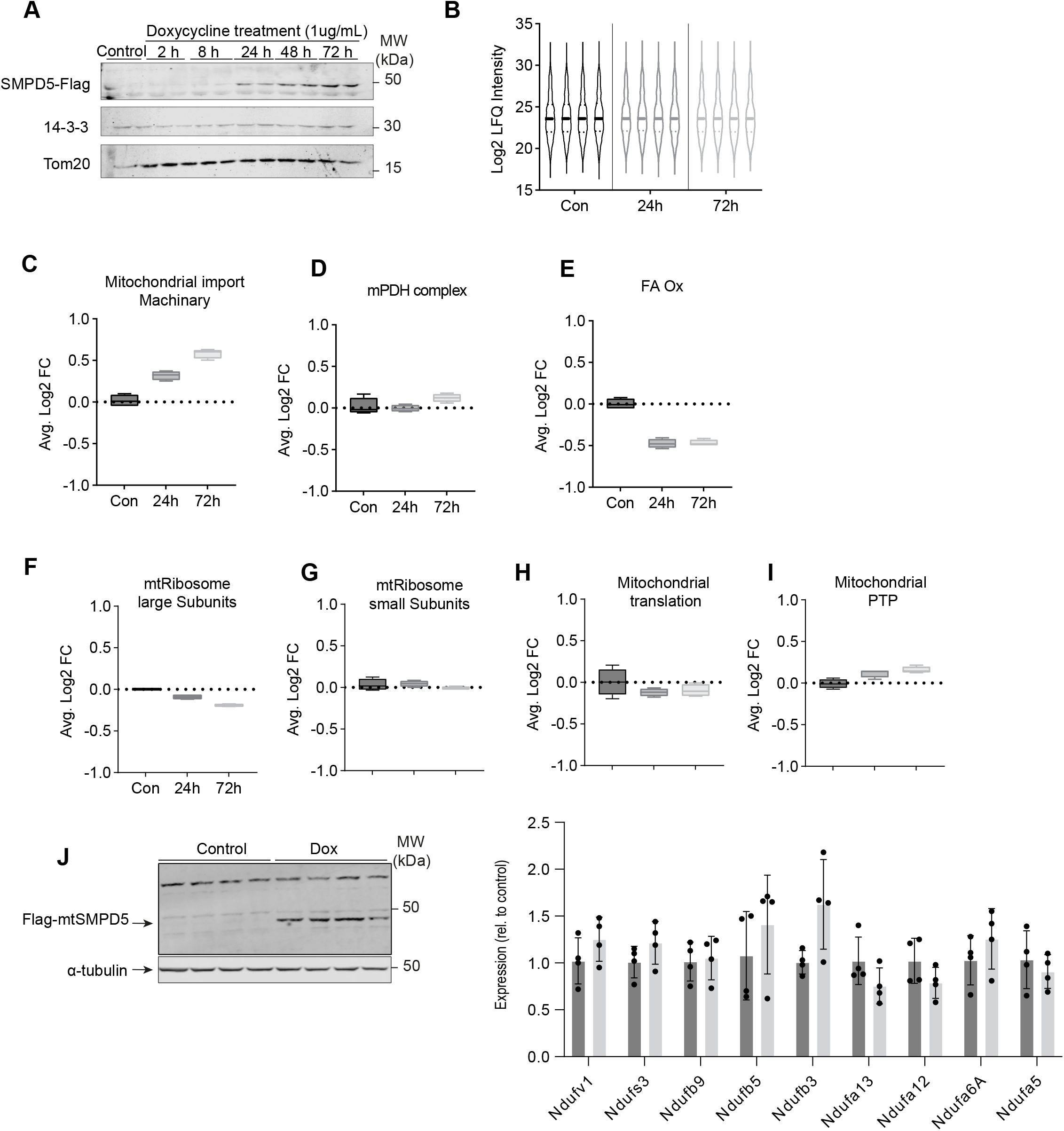
A) L6-myotubes were exposed to doxycycline for different times to promote mtSMPD5 overexpression. The Immunoblot of Flag is shown in A. B) Violin plot of proteomic data after median normalised. C - I) Summed intensities of protein, subunits associated with a specific process as denoted on top of each graph. N = 4 ± S.E.M. J) L6-myotubes were exposed to doxycycline to promote mtSMPD5 overexpression. The Immunoblot of Flag is shown in the left panel. mRNA levels for different complex I subunits were determined by RT-PCR and normalised against beta actin N = 4 ± S.D.

**Figure S7.**
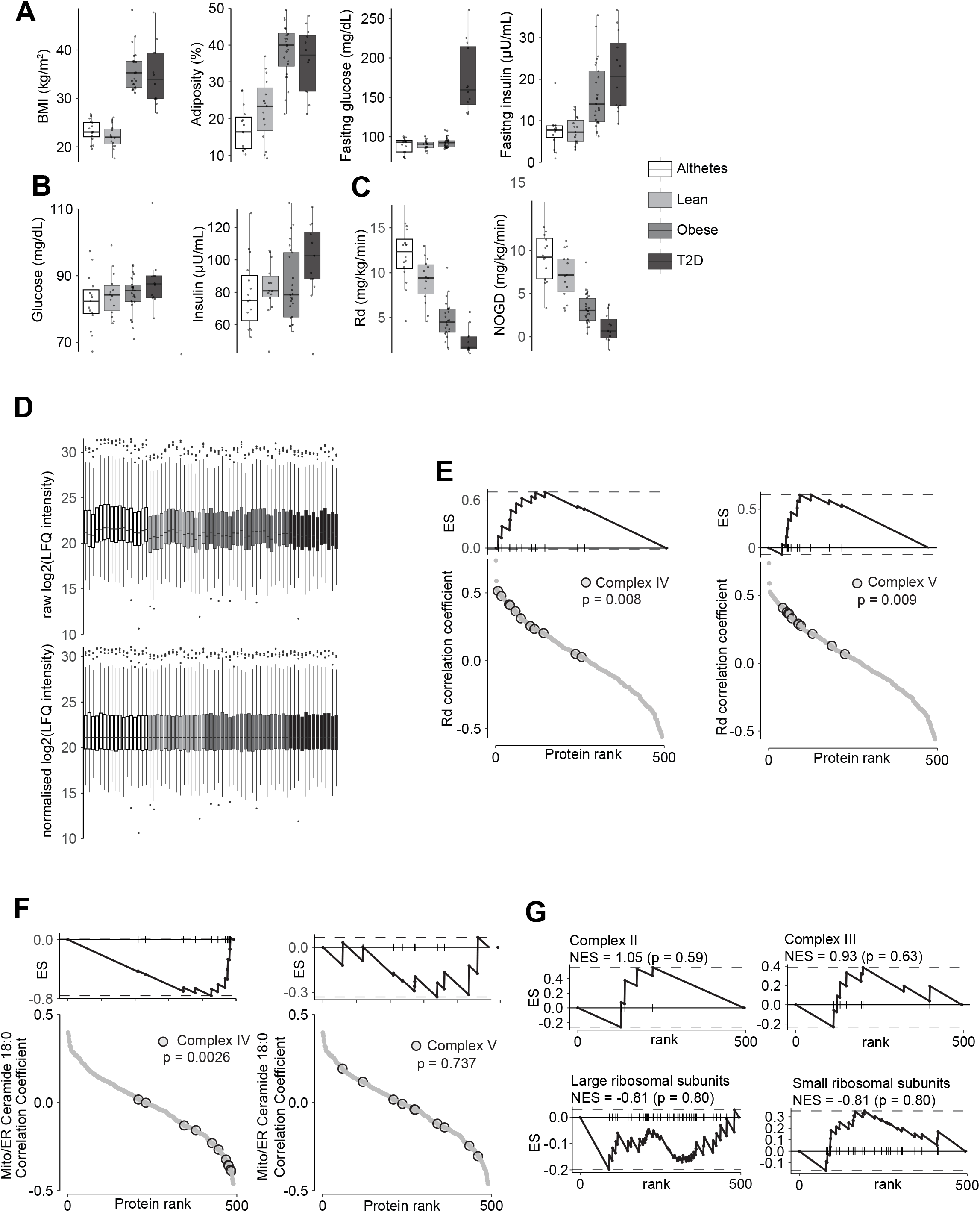
A) Box plot of all proteomics samples before (top) and after (bottom) median normalisation. B) Pairwise comparisons of mitochondrial proteome between all four groups. C) Proteins rank against rate glucose disappearance during clamp (Rd) correlation. Proteins within complex IV and V of the electron transport chain associate with Rd. D) Association of proteins within Complex II, III, small and large ribosomal subunits with Rd.

**Supplementary figure 8.**
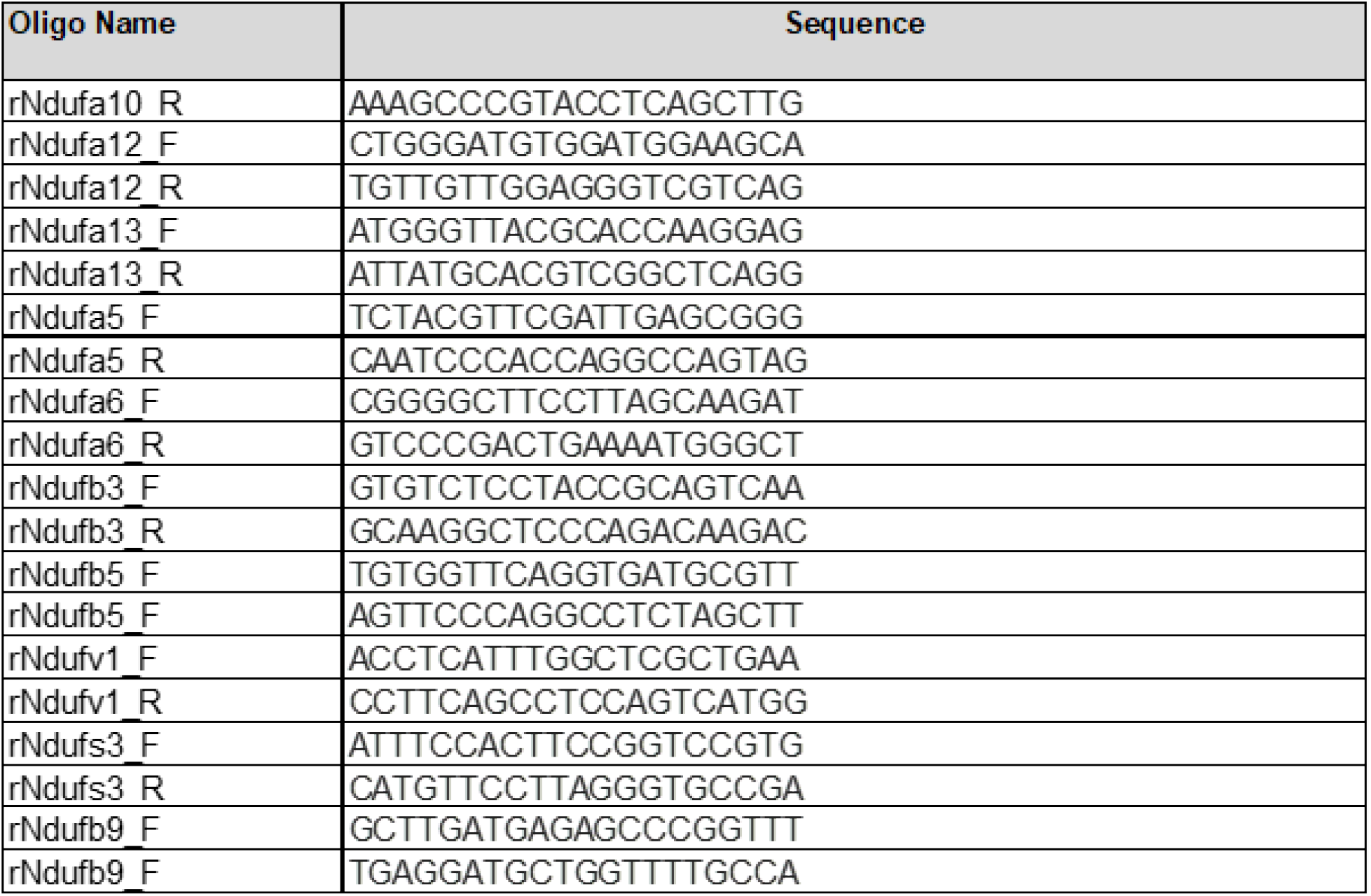

## Lead contact

Further information and requests for resources and reagents should be directed to and will be fulfilled by the Lead Contact, David James or James Burchfield

## Materials availability

This study generated two new molecular tools to overexpress mitochondria targeted SMPD5 and ASAH1. The plasmids are available upon request.

## Data availability

All Lipidomic analyses are available as supplementary information.

## EXPERIMENTAL MODEL

### Administration of P053 to mice

Mice of the C57BL6/J strain were obtained from the Animal Resources Centre of Perth (WA, Australia). Mice were housed in a controlled 12:12 h light-dark cycle, and they had ad libitum access to water and food. The oral gavage administration of P053 (5 mg/kg) was performed daily, while the control animals received vehicle (2% DMSO). The experiments were approved by the UNSW animal care and ethics committee (ACEC 15/48B), and followed guidelines issued by the National Health and Medical Research Council of Australia.

### Cell lines

Mycoplasma-free L6 myotubes overexpressing HA-GLUT4 and HeLa cell lines were used for all in vitro experiments (detailed below each legend). HA-GLUT4 overexpression is essential for studying insulin sensitivity in vitro as we previously described^9^. L6-myoblast and HeLa cells were cultured in Dulbecco’s Modified Eagle Medium (DMEM) (Gibco by Life Technologies) supplemented with 10% foetal bovine serum (FBS) (v/v) (Gibco by Life Technologies) and 2 mM Glutamax (Gibco by Life Technologies) at 37 °C and 10% CO2. L6 myoblasts were differentiated in DMEM/Glutamax/2% Horse serum as previously described^9^. The media was replaced every 48 h for 6 d. For induction of SMPD5 or ASAH1, L6 myotubes were incubated with doxycycline from day 3 until day 6 after the initiation of differentiation. L6-myotubes were used day 7 after the initiation of differentiation. At least 90 % of the cells were differentiated prior to experiments.

## METHOD DETAILS

### Lentiviral transduction

Lentivirus was made by transfecting LentiX-293T (Takara Bio) cells with Lenti-X Packaging Single Shot (Takara Bio) with one of the following plasmids pLVX-Tet3G, pLVX-TRE3G-SMPD5-Myc-DDK or pLVX-TRE3G-ASAH-Myc-DDK according to the manufacturer’s specifications. Virus containing media was collected from the LentiX-293T cells and concentrated using Lenti-X Concentrator (Takara Bio). pLVX-Tet3G virus and polybrene was added to L6-myoblast cells and cells were positively selected using neomycin to create Tet3G expressing cells. The Tet3G expressing L6 cells then were subsequently infected with polybrene and either the pLVX-TRE3G-SMPD5-Myc-DDK or pLVX-TRE3G-ASAH-Myc-DDK virus. Cells were selected using puromycin to create a Tet-inducible SMPD5-Myc-Flag-DDK or ASAH-Myc-Flag-DDK L6 cell line.

### Lipid extraction

Two-phase extraction of lipids from frozen tissue samples (20 mg) or cells was carried out using the methyl-tert-butyl ether (MTBE)/methanol/water (10:3:2.5, v/v/v) method(Matyash et al., 2008). Frozen tissue samples (∼20 mg) were homogenised in 0.2 mL methanol (0.01% butylated hydroxytoluene, BHT) using a Precellys 24 homogenizer and Cryolys cooling unit (Betin Technologies) with CK14 (1.4-mm ceramide) beads. HeLa cells and L6-myotubes were washed with PBS and scraped into 0.6 mL of ice-cold methanol(Turner et al., 2018). Mitochondrial pellets were washed twice to remove BSA from the fraction (see below), and 30 ug of mitochondrial protein was used for extraction. The homogenates were spiked with an internal standard mixture (2 nmole of 18:1/15:0 d5-diacylglycerol and 18:1/17:0 SM, 4 nmole 14:0/14:0/14:0/14:0 cardiolipin, 5 nmole d7-cholesterol, 500 pmole 18:1/17:0 ceramide, and 200 pmole d17:1 sphingosine and d17:1 S1P), then transferred to 10 mL screw cap glass tubes. MTBE (1.7 mL) was added and the samples were sonicated for 30 min in an ice-cold sonicating water bath (Thermoline Scientific, Australia). Phase separation was induced by adding 417 µL of mass spectrometry-grade water with vortexing (max speed for 30 sec), then centrifugation (1000 x g for 10 min). The upper organic phase was transferred into 5 mL glass tubes. The aqueous phase was re-extracted 3 times (MTBE/methanol/water 10:3:2.5), combining the organic phase in the 5 mL glass tube. The organic phase was dried under vacuum in a Savant SC210 SpeedVac (Thermo Scientific). Dried lipids were resuspended in 500 μL of 80% MeOH/ 0.2 % formic acid / 2mM ammonium formate and stored at −20°C until analysis.

### Lipid quantification

Lipids were quantified by selected reaction monitoring on a TSQ Altis triple quadrupole mass spectrometer coupled to a Vanquish HPLC system (ThermoFisher Scientific). Lipids were separated on a 2.1 100 mm Waters Acquity UPLC C18 column (1.7 µM pore size) using a flow rate of 0.28 mL/min. Mobile phase A was 0.1% formic acid, 10 mM ammonium formate in 60% acetonitrile/40% water. Mobile phase B was 0.1% formic acid and 10mM ammonium formate in 90% isopropanol/10% acetonitrile. Total run time was 25 min, starting at 20% B and holding for 3 min, increasing to 100% B from 3-14 min, holding at 100% from 14-20 min, returning to 20% B at 20.5 min, and holding at 20% B for a further 4.5 min. Ceramides, sphingomyelin, sphingosine, and sphingosine 1-phosphate were identified as the [M+H]+ precursor ion, with m/z 262.3 (sphinganine), 264.3 (sphingosine), or 266.3 (sphinganine) product ion, and m/z 184.1 product ion in the case of sphingomyelin. Diacylglycerols (DAGs) were identified as the [M+NH4]+ precursor ion and product ion corresponding to neutral loss of a fatty acid + NH3. Cardiolipins were identified as the [M+H]+ precursor ion and product ion corresponding to neutral loss of a DAG. Cholesterol was detected using precursor m/z 369.4 and product m/z 161.1. TraceFinder software (ThermoFisher) was used for peak alignment and integration. The amount of each lipid was determined relative to its class-specific internal standard. Lipidomic profiling of skeletal muscle tissue was performed exactly as described (Turner et al, 2018).

### Mass spectrometry sample preparation

Isolated mitochondria were defrosted and centrifuged at 4 °C at 4,000 x g for 15 min, and supernatant was removed. The mitochondrial pellet was resuspended in 100 uL 2 % SDC in Tris-HCl buffer (100 mM; pH 8.0) and the protein concentration determined by BCA assay. 10 ug of each sample was aliquoted and volume adjusted to 50 uL with milli-Q water, and samples were reduced and alkylated by addition of TCEP and CAA (10 and 40 mM respectively) at 60 °C for 20 minutes. Once cooled to room temperature, 0.4 ug MS grade trypsin and Lys-C were added to each sample, and proteins were digested overnight (16 h) at 37 °C. Peptides were prepared for MS analysis by SDB-RPS stage tips. 2 layers of SDB-RPS material was packed into 200 µL tips and washed by centrifugation of StageTips at 1,000 x g for 2 min in a 96-well adaptor with 50 µL acetonitrile followed by 0.2% TFA in 30% methanol and then 0.2% TFA in water. 50 µL of samples were loaded to StageTips by centrifugation at 1,000 g for 3 min. Stage tips were washed with subsequent spins at 1,000 g for 3 min with 50 uL 1% TFA in ethyl acetate, then 1% TFA in isopropanol, and 0.2% TFA in 5% ACN. Samples were eluted by addition of 60 µL 60% ACN with 5% NH4OH4. Samples were dried by vacuum centrifugation and reconstituted in 30 µL 0.1% TFA in 2% ACN.

### Mass spectrometry acquisition and analysis

Samples were analysed using a Dionex UltiMate™ 3000 RSLCnano LC coupled to a Exploris Orbitrap mass spectrometer. 3 µL of peptide sample was injected onto an in-house packed 75 μm × 40 cm column (1.9 μm particle size, ReproSil Pur C18-AQ) and separated using a gradient elution, with Buffer A consisting of 0.1 % formic acid in water and Buffer B consisting of 0.1% formic acid in 80% ACN. Samples were loaded to the column at a flow rate 0.5 µL min-1 at 3% B for 14 min, before dropping to 0.3 µL min-1 over 1 min and subsequent ramping to 30% B over 110 min, then to 60% B over 5 min and 98% B over 3 min and held for 6 min, before dropping to 50% and increasing flow rate to 0.5 µL min-1 over 1 min. Eluting peptides were ionised by electrospray with a spray voltage of 2.3 kV and a transfer capillary temperature of 300°C. Mass spectra were collected using a DIA method with varying isolation width windows (widths of m/z 22-589) between 350 - 1650 according to Supplementary Table 1. MS1 spectra were collected between m/z 350 - 1650 m/z at a resolution of 60,000. Ions were fragmented with an HCD collision energy at 30% and MS2 spectra collected between m/z 300-2000 at resolution of 30,000, with an AGC target of 3e5 and the maximum injection time set to automatic. Raw data files were searched using DIA-NN using library generated from a 16-fraction high pH reverse phase library^83^. The protease was set to Trypsin/P with 1 missed cleavage, N-term M excision, carbamidomethylation and M oxidation options on. Peptide length was set to 7-30, precursor range 350-1650, and fragment range 300-2000, and FDR set to 1%.

### Statistical Analysis of L6 mitochondrial proteome

Mouse MitoCarta(REF) was mapped to Rattus norvegicus proteins using OrthoDB identifiers downloaded from Uniprot. The Rat MitoCarta was used to annotate the L6 proteome. Manual scanning of the annotation revealed a number of known mitochondrial proteins not captured using this approach. Proteins were therefore classified as mitochondria if they were annotated by our mouse:rat Mitocarta (380 proteins), contained “mitochondrial” in the protein name (78 additional proteins; 231 overlap with mitocarta) or if the first entry under Uniprot “Subcellular location” was mitochondria (97 additional proteins; 374 overlap with Mitocarta or protein name). LFQ intensities were Log 2 transformed and normalised to the median of the mitochondrially annotated proteins. Identification of differentially regulated proteins was performed using moderated t-tests^84^. Functional enrichment was performed using the STRING web-based platform^85^.

### Statistical Analysis of human proteome and mito-ER lipidome

Analysis of the human proteome and mito-ER lipidome were performed with R (version 4.2.1). Identification of differentially regulated proteins between each group were performed using the R package limma^86^ and p-values were corrected with p.adjust (method = “fdr”) within each comparison. Correlations were calculated with biweight midcorrelations from the R package WGCNA^87^. Gene set enrichment was performed with the R package clusterProfiler^88^ utilising pathways from Reactome for differentially regulated proteins^89^. Custom mitochondrial genes were constructed from HGNC Database^90^ and enrichment and enrichment figures were done with the R package fgsa (https://www.biorxiv.org/content/10.1101/060012v3).

### High pH reverse phase fractionation and library generation

A pooled sample was made by combining 1 uL of each sample and fractionated by high pH reverse phase liquid chromatography. 50 uL of pooled sample was injected onto a Waters XBridge Peptide BEH C18 column (4.6 x 250 mm, 130 Å, 3.5 um) using a ThermoScientific UltiMate 3000 BioRS System and peptides separated using gradient elution at 1 mL min-1, with the column oven set to 30 °C. Buffer A consisted of 10 mM ammonium formate, and Buffer B consisted of 10 mM ammonium formate in 80 % acetonitrile, which both adjusted to pH 9.0 with ammonium hydroxide. Initially Buffer B was set to 10 % and ramped up to 40 % over 11 minutes, before ramping up to 100 % B over 1 minute and held for 5 min before returning to 10 % for re-equilibration. Peptides were separated into 64 fractions collected between 3.45 min to 14.5 min, and samples were concatenated into 16 final fractions. Fractions were dried using a GeneVac 2.0 vacuum centrifuge using the HPLC program, with a max temperature of 60 °C. Fractions were resuspended in 10 uL 0.1% TFA in 2% ACN and 2 uL was injected and separated as described for DIA samples above, however, MS was acquired in a DDA manner. An MS1 was collected between m/z 350-1650 with a resolution of 60,000. The top 15 most intense precursors were selected from fragmentation with an isolation window of 1.4 m/z, resolution of 15,000, HCD collision energy of 30 %, with an exclusion window of 30 s. Raw files were searched with MaxQuant against a FASTA file containing the reviewed UniProt human proteome (downloaded May 2020).

### Matrigel-coated plates

Matrigel diluted 1:100 v/v in ice-cold PBS was dispensed into 96-well plates (Eppendorf Cell Imaging plate, UNSPSC 41122107; and Perkin Elmer Cell Carrier Ultra, Cat# 6055300) and incubated for 2 h at room temperature. Before use, plates were washed twice in PBS at room temperature.

### HA-GLUT4 assay

HA-GLUT4 levels on the plasma membrane were determined as previously described^9,91^. L6-myotubes stably overexpressing HA-Glut4 were washed twice with warm PBS and serum-starved for two hours (in DMEM/0.2% BSA/GlutaMAX/with 220 mM bicarbonate (pH 7.4) at 37 °C, 10 % CO2). Cells were then stimulated with insulin for twenty minutes, after which the cells were placed on ice and washed three times with ice cold PBS. Cells were blocked with ice cold 10 % horse serum in PBS for 20 min, fixed with 4 % paraformaldehyde (PFA) for 5 min on ice and 20 min at room temperature. PFA was quenched with 50 mM glycine in PBS for 5 min at room temperature. We measured the accessibility of the HA epitope to an anti-HA antibody (Covance, 16B12) for 1 h at room temperature. Cells were then incubated with 20 mg/mL goat anti-mouse Alexa-488-conjugated secondary antibody (Thermo Fisher Scientific) for 45 min at room temperature. The determination of total HA-GLUT4 was performed in a separate set of cells following permeabilization with 0.01% saponin (w/v) and anti-HA staining (as above). Each experimental treatment group had its own total HA-GLUT4. A FLuostar Galaxy microplate reader (BMG LABTECH) was used to measure fluorescence (excitation 485 nm/emission 520 nm). Surface HA-GLUT4 was expressed as a fold over control insulin condition.

### Induction of insulin resistance

To promote insulin resistance, cells were stimulated for 16 h with 150 μM palmitate-BSA or EtOH-BSA as control. The palmitate was complexed with BSA as previously described^9^. Briefly, fatty acid was dissolved in 50% ethanol and then diluted 25 times in 10.5 % fatty acid free BSA solution. These stock solutions were further diluted in culture media to reach a final concentration of 150 μM (Final lipid:BSA ratio 4:1).

### Glycogen synthesis assay

L6 myotubes overexpressing ASAH were grown and differentiated in 12-well plates, as described in the Cell lines section, and stimulated for 16 h with palmitate-BSA or EtOH-BSA, as detailed in the Induction of insulin resistance section.

On day seven of differentiation, myotubes were serum starved in plain DMEM for 3 and a half hours. After incubation for 1 hour at 37 °C with 2 µCi/ml D-[U-14C]-glucose in the presence or absence of 100 nM insulin, glycogen synthesis assay was performed, as previously described ^92^

### Coenzyme Q determination

CoQ9 and CoQ10 content in cell lysates and mitochondrial fractions were determined as described previously(Burger et al., 2020). Aliquots of 15 µg mitochondrial protein as prepared below were adjusted to a volume of 100 µL with water and subsequently mixed with 250 μL ice-cold methanol containing 0.1% HCl, 20 µL internal standard (CoQ8, 200 pmol in hexane, Avanti Polar Lipids) and 300 μL of hexane. The mixture was vortexed for 30 sec, centrifuged (9,000 g x 5 min) and the supernatant was transferred into deepwell plate 96/1000 uL (Cat. numb. 951032905). Samples were dried using a rotary evaporator (GeneVac, low BP at 45 °C for 40 min). The resulting dried lipids were re-dissolved in 100 uL of 100 % EtOH (HPLC grade), transferred into HPLC vials and stored at −20 °C until analysis by LC/MS.

LC-MS/MS was performed on a Vanquish LC (ThermoFisher) coupled to a TSQ Altis triple quadrupole mass spectrometer (Thermo Fisher Scientific). Samples were kept at in the autosampler at 4 °C and 15 µL was injected on onto column (50 x 2.1 mm, Kinetex 2.6 μm XB-X18 100 A) at 45 °C, and CoQ8, CoQ9 and CoQ10 were separated by gradient elution using mobile phase A (2.5 mM ammonium formate in 95:5 methanol:isopropanol) and mobile phase B (2.5 mM ammonium formate in 100% isopropanol) at 0.8 mL/min. An initial concentration of 0 % B was held for 1 min before increasing to 45 % B over 1 min and held for 1 min, before decreasing back to 0 % B over 0.5 min and column re-equilibrated over 1.5 min. Under these conditions, CoQ8 eluted at 1.0 min, CoQ9 at 1.6 min and CoQ10 at 2.0 min. Eluent was then directed into the QqQ with a source voltage of 3.5 kV, sheath gas set to 2, auxiliary gas set to 2, and a transfer capillary temperature of 350 °C Ammonium adducts of each of the analytes were detected by SRM with Q1 and Q3 resolution set to 0.7 FWHM with the following parameters: [CoQ8+NH4]+, m/z 744.9 ® 197.1 with collision energy 32.76; [CoQ9+NH4]+, m/z 812.9®197.1 with collision energy 32.76; [CoQ9H2+NH4]+, m/z 814.9®197.1 with collision energy 36.4; and [CoQ10+NH4]+, m/z 880.9 ®197.1 with collision energy 32.76. CoQ9 and CoQ10 areas were normalised to the internal standard CoQ8 levels (20 ng/mL). CoQ9 and CoQ10 were quantified against external standard curves generated from authentic commercial standards obtained from Sigma Aldrich (USA).

### Mitochondrial isolation

Mitochondrial isolation from cultured L6-myotubes and was performed as described elsewhere^93,94^. Briefly, cells were homogenised in an ice-cold mitochondrial isolation buffer (5 mM HEPES, 0.5 mM EGTA, 200 mM mannitol and 0.1 % BSA, pH 7.4 containing protease inhibitors) using a Cell Homogenizer with 18-micron ball. Cells were passed through the Cell Homogenizer 10 times using 1 mL syringe. Cell Homogenizer was equilibrated with 1 mL of ice-cold isolation buffer prior to the experiment. Homogenates were centrifuged at 700 g for 10 min and the supernatant centrifuged at 10,300 g for 10 min to generate the crude mitochondrial pellet. The 10,300 g pellet was resuspended in 1 mL of isolation buffer and transferred into a polycarbonate tube containing 7.9 mL of 18% Percoll in the homogenization buffer and centrifuged at 95,000 g at 4 °C for 30 min. The mitochondrial pellet was collected and diluted in a homogenization buffer (1 mL) and centrifuged at 10,000 g for 10 min at 4 °C. The supernatant was discarded, and the pellet was washed with a homogenization buffer without BSA followed by protein quantification with BCA protein assay.

Mitochondria from adult skeletal muscle (from mixed hindlimb muscle) were isolated by differential centrifugation as described previously^95^. Briefly, muscle was diced in CP-1 medium (100 mM KCl, 50 mM Tris/HCl, pH 7.4, and 2 mM EGTA), digested on ice for 3 min in CP-2 medium [CP-1, to which was added 0.5% (w/v) BSA, 5 mM MgCl2, 1 mM ATP and 2.45 units ml–1 Protease Type VIII (Sigma P 5380)] and homogenised using an ultra-turrax homogenizer. The homogenate was spun for 5 min at 500 g and 4°C. The resulting supernatant was subjected to a high-speed spin (10,600 g, 10 min, 4°C) and the mitochondrial pellet was resuspended in CP-1. The 10,600 g spin cycle was repeated, the supernatant removed and the mitochondrial pellet snapped frozen.

### Western Blotting

After insulin stimulation or mitochondrial isolation, samples were tip sonicated in 2% SDS-RIPA. Insoluble material was removed by centrifugation at 21,000 g x 10 min. Protein concentration was determined by bicinchoninic acid method (Thermo Scientific). 10 μg of protein was resolved by SDS-PAGE and transferred to PDVF membranes. Membranes were blocked in Tris-buffered saline (TBS) 4 % skim milk for 30 min at room temperature, followed by primary antibody incubation (detailed antibody is provided in “Key Resource Table”). Membranes were washed in TBS 0.1% tween (TBS-T) and incubated with appropriate secondary antibodies (IRDye700- or IRDye800-conjugated) in TBS-T 2% skim milk for 45 min at room temperature. Images were obtained by using 700- or 800-nm channels using Odyssey IR imager. Densitometry analysis of immunoblots was performed using Image Studio Lite (version 5.2). Uncropped Western blots are provided in Supplementary Material.

### Seahorse extracellular flux analyses

Mitochondrial respiration (*J*O_2_) of intact cells were measured using an XF HS mini Analyser Extracellular Flux Analyzer (Seahorse Bioscience, Copenhagen, Denmark). L6 myoblasts were seeded and differentiated in Seahorse XFp culture plates coated with matrigel and assayed after incubation at 37°C without CO_2_ for 1 hour. Prior to the assay, cells were washed 3 times with PBS, once with bicarbonate-free DMEM buffered with 30 mM Na-HEPES, pH 7.4 (DMEM/HEPES), and then incubated in DMEM/HEPES supplemented with 0.2% (w/v) BSA, 25 mM glucose, 1 mM GlutaMAX and 1 mM glutamine (Media B), for 1.5 h in a non-CO2 incubator at 37 °C. During the assay, respiration was assayed with mix/wait/read cycles of 2/0/2 min for L6 myotubes. Following assessment of basal respiration, the following compounds (final concentrations in parentheses) were injected sequentially: oligoymcin (10 μg/ml), BAM15 (10 mM), rotenone/antimycin A (5 μM / 10 μM). All of these reagents were obtained from Sigma-Aldrich. basal (baseline - Ant./Rot), ATP-linked respiration (determined by basal – oligomycin), maximal respiration (calculated by FCCP – AntA/Rot) and non-mitochondrial respiration (equal to AntA/Rot) was determined as previously described^96^. Protein concentration was determined immediately after the assy and data are presented as O2/min. Complex specific activity in permeabilized cells were performed according to^97^. The cells were seeded and the media was changed to a buffer consisting of 70 mM sucrose, 220 mM Mannitol, 10 mM KH2PO4, 5 mM MgCl2, 2 mM Hepes (pH 7.2), 1 mM EGTA, and 0.4% BSA. Then, flux measurements began after taking three baseline measurements. The cells were permeabilized by adding digitonin (1 nM) and 1 mM ADP, followed by injecting respiratory complex substrates or ADP only (complex I, glutamate/malate (5 mM/2.5 mM); complex II, succinate/rotenone (10 mM/1 μM); complex III, and complex IV, N,N,N,N-tetramethyl-p-phenylenediamine/ascorbate (0.5 mM/2 mM)). Subsequently, oligomycin (1 μg/ml) and respective complex inhibitors were added (complex I, 1 μM rotenone; complexes II and III, 20 μM antimycin A; complex IV, 20 mM sodium azide). Wells where cells detached from the plate during the assay were excluded from the analysis.

### Mitochondrial membrane potential

Mitochondrial membrane potential was measured by loading cells with 20nM tetramethylrhodamine, ethyl ester (TMRM+, Life Technologies) for 30 min at 37◦C plus mitotracker deep Red (MTDR). MTDR was used to normalize the fluorescence among the different mitochondrial populations as previously reported^96^. TMRM+ fluorescence was detected using the excitation-emission λ545–580/590 nm and MTDR was detected using an ex/em ~644/665 nm using confocal microscopy. The mitochondrial membrane potential was evaluated as raw fluorescence intensity of background-corrected images.

### Mitochondrial oxidative stress

MitoSOX Red was administered as described by the manufacturer (Molecular Probes); At the end of the induction period cells were washed twice with PBS and incubated with 0.5 μM MitoSOX Red for 30 min plus mitotracker deep Red (MTDR). MTDR was used to normalize the fluorescence among the different mitochondrial populations as previously reported. Cells were cultured in low-absorbance, black-walled 96-well plates. After MitoSOX treatment cells were quickly washed with PBS and fluorescence was detected on a confocal microscope. MitoSOX fluorescence was detected using the excitation-emission λ396/610 nm and MTDR was detected using an ex/em ~644/665 nm using confocal microscopy.

### RT-PCR

RNA was extracted by addition of TRIzol (Thermo-fisher) followed by addition of 0.1x volume of 1-bromo-3-chloropropane. Samples were centrifuged at 13,000xg for 15 minutes for phase separation. The clear phase was transferred to a fresh tube and an equal volume of isopropanol was added. Samples were centrifuged at 13,000xg for 10 minutes to precipitate RNA. RNA was washed three times in 70% ethanol with centrifugation. RNA was resuspended in DEPC treated water and quantified on a NanoDrop 2000 (Thermo Scientific). RNA was reverse transcribed to cDNA using PrimeScript Reverse Transcriptase(Takara) as per the manufacturer’s instructions. qPCR was performed using FastStart SYBR Green MasterMix (2x) (Biorad) as per the manufacturer’s instructions. All primers sequence van be found in sup. Fig. 8

### Statistical analysis

Data are presented as mean ± S.E.M. Statistical tests were performed using GraphPad Prism version 9. HA-GLUT4 and Western blot assay was analysed by using Kruskal-Wallis with Dunn’s multiple comparisons test. CoQ and ceramide abundance were analysed with ordinary one-Way ANOVA and Dunnett’s multiple comparison test. Finally, for comparison of two groups (CoQ and Ceramides levels in mice) we use Student’s *t*-test. Significant effects were defined as p<0.05 by these tests as reported in the Figures.

## References

1 Hill MM, Clark SF, Tucker DF, Birnbaum MJ, James DE, Macaulay SL. A role for protein kinase Bbeta/Akt2 in insulin-stimulated GLUT4 translocation in adipocytes. Mol Cell Biol 1999; 19: 7771–7781.

2 Cong LN, Chen H, Li Y, Zhou L, McGibbon MA, Taylor SI et al. Physiological role of Akt in insulin-stimulated translocation of GLUT4 in transfected rat adipose cells. Mol Endocrinol 1997; 11: 1881–1890.

3 James DE, Stöckli J, Birnbaum MJ. The aetiology and molecular landscape of insulin resistance. Nat Rev Mol Cell Biol 2021; 22: 751–771.

4 Riehle C, Abel ED. Insulin Signaling and Heart Failure. Circ Res 2016; 118: 1151–1169.

5 Leitner BP, Siebel S, Akingbesote ND, Zhang X, Perry RJ. Insulin and cancer: a tangled web. Biochem J 2022; 479: 583–607.

6 Fazakerley DJ, Chaudhuri R, Yang P, Maghzal GJ, Thomas KC, Krycer JR et al. Mitochondrial CoQ deficiency is a common driver of mitochondrial oxidants and insulin resistance. Elife 2018; 7. doi:10.7554/eLife.32111.

7 Holland WL, Summers SA. Sphingolipids, insulin resistance, and metabolic disease: new insights from in vivo manipulation of sphingolipid metabolism. Endocr Rev 2008; 29: 381–402.

8 Anderson EJ, Lustig ME, Boyle KE, Woodlief TL, Kane DA, Lin C-T et al. Mitochondrial H2O2 emission and cellular redox state link excess fat intake to insulin resistance in both rodents and humans. J Clin Invest 2009; 119: 573–581.

9 Hoehn KL, Salmon AB, Hohnen-Behrens C, Turner N, Hoy AJ, Maghzal GJ et al. Insulin resistance is a cellular antioxidant defense mechanism. Proc Natl Acad Sci U S A 2009; 106: 17787–17792.

10 Fazakerley DJ, Minard AY, Krycer JR, Thomas KC, Stöckli J, Harney DJ et al. Mitochondrial oxidative stress causes insulin resistance without disrupting oxidative phosphorylation. J Biol Chem 2018; 293: 7315–7328.

11 Hatefi Y, Haavik AG, Fowler LR, Griffiths DE. Studies on the electron transfer system. XLII. Reconstitution of the electron transfer system. J Biol Chem 1962; 237: 2661–2669.

12 Frerman FE. Acyl-CoA dehydrogenases, electron transfer flavoprotein and electron transfer flavoprotein dehydrogenase. Biochem Soc Trans 1988; 16: 416–418.

13 Jones ME. Pyrimidine nucleotide biosynthesis in animals: genes, enzymes, and regulation of UMP biosynthesis. Annu Rev Biochem 1980; 49: 253–279.

14 Montini G, Malaventura C, Salviati L. Early coenzyme Q10 supplementation in primary coenzyme Q10 deficiency. N Engl J Med 2008; 358: 2849–2850.

15 Kühl I, Miranda M, Atanassov I, Kuznetsova I, Hinze Y, Mourier A et al. Transcriptomic and proteomic landscape of mitochondrial dysfunction reveals secondary coenzyme Q deficiency in mammals. Elife 2017; 6. doi:10.7554/eLife.30952.

16 Calvo E, Cogliati S, Hernansanz-Agustín P, Loureiro-López M, Guarás A, Casuso RA et al. Functional role of respiratory supercomplexes in mice: SCAF1 relevance and segmentation of the Qpool. Sci Adv 2020; 6: eaba7509.

17 Ates O, Bilen H, Keles S, Alp HH, Keleş MS, Yıldırım K et al. Plasma coenzyme Q10 levels in type 2 diabetic patients with retinopathy. Int J Ophthalmol 2013; 6: 675–679.

18 El-ghoroury EA, Raslan HM, Badawy EA, El-Saaid GS, Agybi MH, Siam I et al. Malondialdehyde and coenzyme Q10 in platelets and serum in type 2 diabetes mellitus: correlation with glycemic control. Blood Coagul Fibrinolysis 2009; 20: 248–251.

19 Zozina VI, Covantev S, Goroshko OA, Krasnykh LM, Kukes VG. Coenzyme Q10 in Cardiovascular and Metabolic Diseases: Current State of the Problem. Curr Cardiol Rev 2018; 14: 164–174.

20 Barcelos IP de, Haas RH. CoQ10 and Aging. Biology 2019; 8. doi:10.3390/biology8020028.

21 Højlund K, Yi Z, Lefort N, Langlais P, Bowen B, Levin K et al. Human ATP synthase beta is phosphorylated at multiple sites and shows abnormal phosphorylation at specific sites in insulin-resistant muscle. Diabetologia 2010; 53: 541–551.

22 Diaz-Vegas A, Sanchez-Aguilera P, Krycer JR, Morales PE, Monsalves-Alvarez M, Cifuentes M et al. Is Mitochondrial Dysfunction a Common Root of Noncommunicable Chronic Diseases? Endocr Rev 2020; 41. doi:10.1210/endrev/bnaa005.

23 Chaurasia B, Summers SA. Ceramides in Metabolism: Key Lipotoxic Players. Annu Rev Physiol 2021; 83: 303–330.

24 Summers SA, Garza LA, Zhou H, Birnbaum MJ. Regulation of insulin-stimulated glucose transporter GLUT4 translocation and Akt kinase activity by ceramide. Mol Cell Biol 1998; 18: 5457–5464.

25 Schubert KM, Scheid MP, Duronio V. Ceramide inhibits protein kinase B/Akt by promoting dephosphorylation of serine 473. J Biol Chem 2000; 275: 13330–13335.

26 Powell DJ, Hajduch E, Kular G, Hundal HS. Ceramide disables 3-phosphoinositide binding to the pleckstrin homology domain of protein kinase B (PKB)/Akt by a PKCzeta-dependent mechanism. Mol Cell Biol 2003; 23: 7794–7808.

27 Kono T, Barham FW. The relationship between the insulin-binding capacity of fat cells and the cellular response to insulin. Studies with intact and trypsin-treated fat cells. J Biol Chem 1971; 246: 6210–6216.

28 Hoehn KL, Hohnen-Behrens C, Cederberg A, Wu LE, Turner N, Yuasa T et al. IRS1-independent defects define major nodes of insulin resistance. Cell Metab 2008; 7: 421–433.

29 Di Paola M, Cocco T, Lorusso M. Ceramide interaction with the respiratory chain of heart mitochondria. Biochemistry 2000; 39: 6660–6668.

30 Perreault L, Newsom SA, Strauss A, Kerege A, Kahn DE, Harrison KA et al. Intracellular localization of diacylglycerols and sphingolipids influences insulin sensitivity and mitochondrial function in human skeletal muscle. JCI Insight 2018; 3. doi:10.1172/jci.insight.96805.

31 Hammerschmidt P, Ostkotte D, Nolte H, Gerl MJ, Jais A, Brunner HL et al. CerS6-Derived Sphingolipids Interact with Mff and Promote Mitochondrial Fragmentation in Obesity. Cell 2019; 177: 1536–1552.e23.

32 Ertunc ME, Hotamisligil GS. Lipid signaling and lipotoxicity in metaflammation: indications for metabolic disease pathogenesis and treatment. J Lipid Res 2016; 57: 2099–2114.

33 Forsman U, Sjöberg M, Turunen M, Sindelar PJ. 4-Nitrobenzoate inhibits coenzyme Q biosynthesis in mammalian cell cultures. Nat Chem Biol 2010; 6: 515–517.

34 Fernández-Ayala DJ, Martín SF, Barroso MP, Gómez-Díaz C, Villalba JM, Rodríguez-Aguilera JC et al. Coenzyme Q protects cells against serum withdrawal-induced apoptosis by inhibition of ceramide release and caspase-3 activation. Antioxid Redox Signal 2000; 2: 263–275.

35 Koves TR, Ussher JR, Noland RC, Slentz D, Mosedale M, Ilkayeva O et al. Mitochondrial overload and incomplete fatty acid oxidation contribute to skeletal muscle insulin resistance. Cell Metab 2008; 7: 45–56.

36 Bruce CR, Hoy AJ, Turner N, Watt MJ, Allen TL, Carpenter K et al. Overexpression of carnitine palmitoyltransferase-1 in skeletal muscle is sufficient to enhance fatty acid oxidation and improve high-fat diet-induced insulin resistance. Diabetes 2009; 58: 550–558.

37 Sebastián D, Herrero L, Serra D, Asins G, Hegardt FG. CPT I overexpression protects L6E9 muscle cells from fatty acid-induced insulin resistance. Am J Physiol Endocrinol Metab 2007; 292: E677–86.

38 Perdomo G, Commerford SR, Richard A-MT, Adams SH, Corkey BE, O’Doherty RM et al. Increased beta-oxidation in muscle cells enhances insulin-stimulated glucose metabolism and protects against fatty acid-induced insulin resistance despite intramyocellular lipid accumulation. J Biol Chem 2004; 279: 27177–27186.

39 Fisher-Wellman KH, Hagen JT, Neufer PD, Kassai M, Cabot MC. On the nature of ceramide-mitochondria interactions - Dissection using comprehensive mitochondrial phenotyping. Cell Signal 2021; 78: 109838.

40 Das AT, Tenenbaum L, Berkhout B. Tet-On Systems For Doxycycline-inducible Gene Expression. Curr Gene Ther 2016; 16: 156–167.

41 Wu BX, Rajagopalan V, Roddy PL, Clarke CJ, Hannun YA. Identification and characterization of murine mitochondria-associated neutral sphingomyelinase (MA-nSMase), the mammalian sphingomyelin phosphodiesterase 5. J Biol Chem 2010; 285: 17993–18002.

42 Bienias K, Fiedorowicz A, Sadowska A, Prokopiuk S, Car H. Regulation of sphingomyelin metabolism. Pharmacol Rep 2016; 68: 570–581.

43 Feng S, Harayama T, Montessuit S, David FP, Winssinger N, Martinou J-C et al. Mitochondria-specific photoactivation to monitor local sphingosine metabolism and function. Elife 2018; 7. doi:10.7554/eLife.34555.

44 Li CM, Park JH, He X, Levy B, Chen F, Arai K et al. The human acid ceramidase gene (ASAH): structure, chromosomal location, mutation analysis, and expression. Genomics 1999; 62: 223–231.

45 Turner N, Lim XY, Toop HD, Osborne B, Brandon AE, Taylor EN et al. A selective inhibitor of ceramide synthase 1 reveals a novel role in fat metabolism. Nat Commun 2018; 9: 3165.

46 Rath S, Sharma R, Gupta R, Ast T, Chan C, Durham TJ et al. MitoCarta3.0: an updated mitochondrial proteome now with sub-organelle localization and pathway annotations. Nucleic Acids Res 2021; 49: D1541–D1547.

47 Letts JA, Sazanov LA. Clarifying the supercomplex: the higher-order organization of the mitochondrial electron transport chain. Nat Struct Mol Biol 2017; 24: 800–808.

48 Zhu J, Vinothkumar KR, Hirst J. Structure of mammalian respiratory complex I. Nature 2016; 536: 354–358.

49 Brand MD, Nicholls DG. Assessing mitochondrial dysfunction in cells. Biochem J 2011; 435: 297–312.

50 Krycer JR, Elkington SD, Diaz-Vegas A, Cooke KC, Burchfield JG, Fisher-Wellman KH et al. Mitochondrial oxidants, but not respiration, are sensitive to glucose in adipocytes. J Biol Chem 2020; 295: 99–110.

51 Sánchez-Aguilera P, Diaz-Vegas A, Campos C, Quinteros-Waltemath O, Cerda-Kohler H, Barrientos G et al. Role of ABCA1 on membrane cholesterol content, insulin-dependent Akt phosphorylation and glucose uptake in adult skeletal muscle fibers from mice. Biochim Biophys Acta Mol Cell Biol Lipids 2018; 1863: 1469–1477.

52 Rosales-Soto G, Diaz-Vegas A, Casas M, Contreras-Ferrat A, Jaimovich E. Fibroblast growth factor-21 potentiates glucose transport in skeletal muscle fibers. J Mol Endocrinol 2020. doi:10.1530/JME-19-0210.

53 Valladares D, Utreras-Mendoza Y, Campos C, Morales C, Diaz-Vegas A, Contreras-Ferrat A et al. IP3 receptor blockade restores autophagy and mitochondrial function in skeletal muscle fibers of dystrophic mice. Biochim Biophys Acta Mol Basis Dis 2018; 1864: 3685–3695.

54 Chalfant CE, Rathman K, Pinkerman RL, Wood RE, Obeid LM, Ogretmen B et al. De novo ceramide regulates the alternative splicing of caspase 9 and Bcl-x in A549 lung adenocarcinoma cells. Dependence on protein phosphatase-1. J Biol Chem 2002; 277: 12587–12595.

55 Tesfay L, Paul BT, Konstorum A, Deng Z, Cox AO, Lee J et al. Stearoyl-CoA Desaturase 1 Protects Ovarian Cancer Cells from Ferroptotic Cell Death. Cancer Res. 2019; 79: 5355–5366.

56 Morad SAF, Levin JC, Shanmugavelandy SS, Kester M, Fabrias G, Bedia C et al. Ceramide--antiestrogen nanoliposomal combinations--novel impact of hormonal therapy in hormone-insensitive breast cancer. Mol Cancer Ther 2012; 11: 2352–2361.

57 Schägger H, Pfeiffer K. Supercomplexes in the respiratory chains of yeast and mammalian mitochondria. EMBO J 2000; 19: 1777–1783.

58 Lapuente-Brun E, Moreno-Loshuertos R, Acín-Pérez R, Latorre-Pellicer A, Colás C, Balsa E et al. Supercomplex assembly determines electron flux in the mitochondrial electron transport chain. Science 2013; 340: 1567–1570.

59 Maranzana E, Barbero G, Falasca AI, Lenaz G, Genova ML. Mitochondrial respiratory supercomplex association limits production of reactive oxygen species from complex I. Antioxid Redox Signal 2013; 19: 1469–1480.

60 Guarás A, Perales-Clemente E, Calvo E, Acín-Pérez R, Loureiro-Lopez M, Pujol C et al. The CoQH2/CoQ Ratio Serves as a Sensor of Respiratory Chain Efficiency. Cell Rep 2016; 15: 197–209.

61 Gudz TI, Tserng KY, Hoppel CL. Direct inhibition of mitochondrial respiratory chain complex III by cell-permeable ceramide. J Biol Chem 1997; 272: 24154–24158.

62 Pinto SN, Silva LC, Futerman AH, Prieto M. Effect of ceramide structure on membrane biophysical properties: the role of acyl chain length and unsaturation. Biochim Biophys Acta 2011; 1808: 2753–2760.

63 Wu M, Gu J, Guo R, Huang Y, Yang M. Structure of Mammalian Respiratory Supercomplex I1III2IV1. Cell 2016; 167: 1598–1609.e10.

64 Mileykovskaya E, Dowhan W. Cardiolipin-dependent formation of mitochondrial respiratory supercomplexes. Chem Phys Lipids 2014; 179: 42–48.

65 Babiychuk EB, Atanassoff AP, Monastyrskaya K, Brandenberger C, Studer D, Allemann C et al. The targeting of plasmalemmal ceramide to mitochondria during apoptosis. PLoS One 2011; 6: e23706.

66 Babiychuk EB, Monastyrskaya K, Draeger A. Fluorescent annexin A1 reveals dynamics of ceramide platforms in living cells. Traffic 2008; 9: 1757–1775.

67 Bionda C, Portoukalian J, Schmitt D, Rodriguez-Lafrasse C, Ardail D. Subcellular compartmentalization of ceramide metabolism: MAM (mitochondria-associated membrane) and/or mitochondria? Biochem J 2004; 382: 527–533.

68 Novgorodov SA, Wu BX, Gudz TI, Bielawski J, Ovchinnikova TV, Hannun YA et al. Novel pathway of ceramide production in mitochondria: thioesterase and neutral ceramidase produce ceramide from sphingosine and acyl-CoA. J Biol Chem 2011; 286: 25352–25362.

69 Birbes H, El Bawab S, Hannun YA, Obeid LM. Selective hydrolysis of a mitochondrial pool of sphingomyelin induces apoptosis. FASEB J 2001; 15: 2669–2679.

70 Yu J, Novgorodov SA, Chudakova D, Zhu H, Bielawska A, Bielawski J et al. JNK3 signaling pathway activates ceramide synthase leading to mitochondrial dysfunction. J Biol Chem 2007; 282: 25940–25949.

71 García-Ruiz C, Colell A, Marí M, Morales A, Fernández-Checa JC. Direct effect of ceramide on the mitochondrial electron transport chain leads to generation of reactive oxygen species. Role of mitochondrial glutathione. J Biol Chem 1997; 272: 11369–11377.

72 Turpin-Nolan SM, Hammerschmidt P, Chen W, Jais A, Timper K, Awazawa M et al. CerS1-Derived C18:0 Ceramide in Skeletal Muscle Promotes Obesity-Induced Insulin Resistance. Cell Rep 2019; 26: 1–10.e7.

73 Oleinik N, Kim J, Roth BM, Selvam SP, Gooz M, Johnson RH et al. Mitochondrial protein import is regulated by p17/PERMIT to mediate lipid metabolism and cellular stress. Sci Adv 2019; 5: eaax1978.

74 Bonnard C, Durand A, Peyrol S, Chanseaume E, Chauvin M-A, Morio B et al. Mitochondrial dysfunction results from oxidative stress in the skeletal muscle of diet-induced insulin-resistant mice. J Clin Invest 2008; 118: 789–800.

75 Picard M, Shirihai OS. Mitochondrial signal transduction. Cell Metab 2022; 34: 1620–1653.

76 Zorov DB, Juhaszova M, Sollott SJ. Mitochondrial reactive oxygen species (ROS) and ROS-induced ROS release. Physiol Rev 2014; 94: 909–950.

77 Siskind LJ, Kolesnick RN, Colombini M. Ceramide channels increase the permeability of the mitochondrial outer membrane to small proteins. J Biol Chem 2002; 277: 26796–26803.

78 Taddeo EP, Laker RC, Breen DS, Akhtar YN, Kenwood BM, Liao JA et al. Opening of the mitochondrial permeability transition pore links mitochondrial dysfunction to insulin resistance in skeletal muscle. Mol Metab 2014; 3: 124–134.

79 Cho J, Zhang Y, Park S-Y, Joseph A-M, Han C, Park H-J et al. Mitochondrial ATP transporter depletion protects mice against liver steatosis and insulin resistance. Nat Commun 2017; 8: 14477.

80 Walter L, Nogueira V, Leverve X, Heitz MP, Bernardi P, Fontaine E. Three classes of ubiquinone analogs regulate the mitochondrial permeability transition pore through a common site. J Biol Chem 2000; 275: 29521–29527.

81 Galgani JE, Fernández-Verdejo R. Pathophysiological role of metabolic flexibility on metabolic health. Obes Rev 2021; 22: e13131.

82 Hernansanz-Agustín P, Enríquez JA. Functional segmentation of CoQ and cyt c pools by respiratory complex superassembly. Free Radic Biol Med 2021; 167: 232–242.

83 Yang F, Shen Y, Camp DG 2nd, Smith RD. High-pH reversed-phase chromatography with fraction concatenation for 2D proteomic analysis. Expert Rev Proteomics 2012; 9: 129–134.

84 Smyth GK. Linear models and empirical bayes methods for assessing differential expression in microarray experiments. Stat Appl Genet Mol Biol 2004; 3: Article3.

85 Szklarczyk D, Gable AL, Lyon D, Junge A, Wyder S, Huerta-Cepas J et al. STRING v11: protein-protein association networks with increased coverage, supporting functional discovery in genome-wide experimental datasets. Nucleic Acids Res 2019; 47: D607–D613.

86 Ritchie ME, Phipson B, Wu D, Hu Y, Law CW, Shi W et al. limma powers differential expression analyses for RNA-sequencing and microarray studies. Nucleic Acids Res 2015; 43: e47.

87 Langfelder P, Horvath S. WGCNA: an R package for weighted correlation network analysis. BMC Bioinformatics 2008; 9: 559.

88 Wu T, Hu E, Xu S, Chen M, Guo P, Dai Z et al. clusterProfiler 4.0: A universal enrichment tool for interpreting omics data. Innovation (Camb) 2021; 2: 100141.

89 Fabregat A, Jupe S, Matthews L, Sidiropoulos K, Gillespie M, Garapati P et al. The Reactome Pathway Knowledgebase. Nucleic Acids Res 2018; 46: D649–D655.

90 Seal RL, Braschi B, Gray K, Jones TEM, Tweedie S, Haim-Vilmovsky L et al. Genenames.org: the HGNC resources in 2023. Nucleic Acids Res 2023; 51: D1003–D1009.

91 Govers R, Coster ACF, James DE. Insulin increases cell surface GLUT4 levels by dose dependently discharging GLUT4 into a cell surface recycling pathway. Mol Cell Biol 2004; 24: 6456–6466.

92 Zarini S, Brozinick JT, Zemski Berry KA, Garfield A, Perreault L, Kerege A et al. Serum dihydroceramides correlate with insulin sensitivity in humans and decrease insulin sensitivity in vitro. J Lipid Res 2022; 63: 100270.

93 Bui M, Gilady SY, Fitzsimmons REB, Benson MD, Lynes EM, Gesson K et al. Rab32 modulates apoptosis onset and mitochondria-associated membrane (MAM) properties. J Biol Chem 2010; 285: 31590–31602.

94 Frezza C, Cipolat S, Scorrano L. Organelle isolation: functional mitochondria from mouse liver, muscle and cultured fibroblasts. Nat Protoc 2007; 2: 287–295.

95 Montgomery MK, Osborne B, Brandon AE, O’Reilly L, Fiveash CE, Brown SHJ et al. Regulation of mitochondrial metabolism in murine skeletal muscle by the medium-chain fatty acid receptor Gpr84. FASEB J 2019; 33: 12264–12276.

96 Díaz-Vegas AR, Cordova A, Valladares D, Llanos P, Hidalgo C, Gherardi G et al. Mitochondrial Calcium Increase Induced by RyR1 and IP3R Channel Activation After Membrane Depolarization Regulates Skeletal Muscle Metabolism. Front Physiol 2018; 9: 791.

97 Kory N, Uit de Bos J, van der Rijt S, Jankovic N, Güra M, Arp N et al. MCART1/SLC25A51 is required for mitochondrial NAD transport. Sci Adv 2020; 6. doi:10.1126/sciadv.abe5310.

